# Floral scent diversity and evolution in deceptive *Cypripedium,* and post-pollination effects on floral scent and its biosynthesis in *C. henryi*

**DOI:** 10.64898/2026.11.04.760698

**Authors:** Jean-Baptiste Chazalon, Simon Pfanzelt, Christian Siadjeu, Emy Yue Hu, Suvrat Kotagal, Stefan Dötterl, Gudrun Kadereit

## Abstract

**Background and Aims:** Floral scent is a primary driver of pollinator attraction and reproductive isolation in deceptive orchids. Yet, little is known about the variation and evolution of scent in a phylogenetic framework and how pollination affects the biosynthesis and emission of scents in these lineages. This study characterizes scent diversity across *Cypripedium,* a genus of deceptive orchids, examines phylogenetic constraints, and investigates post-pollination effects on scent composition in one of the species.

**Methods:** Floral volatile organic compounds (VOCs) were sampled from 31 *Cypripedium* taxa using dynamic headspace techniques, and analysed by gas chromatography coupled to mass spectrometry (GC/MS). Uni-and multivariate methods were used to analyse intra-and interspecific scent variation and test for phylogenetic signal in floral scent. Post-pollination effects on scent composition were experimentally tested in *C. henryi* using four pollination treatments, combined with RNA-seq analyses of floral tissues to examine transcriptional responses in VOC biosynthetic pathways.

**Key results:** We detected 258 floral VOCs, revealing a high chemical diversity dominated by terpenoids, aliphatics, and aromatic compounds. Floral scent composition was highly species-specific, and closely related taxa often shared only few compounds. Scent composition differed among pollination groups, but these groups explained only a limited proportion of overall scent variation. We detected a significant but weak phylogenetic signal, suggesting limited evolutionary constraint in floral scent. Pollination had limited effect on *C. henryi’*s floral scent and volatile biosynthesis gene expression, despite a significant decrease in total scent emission after pollinium deposition. Transcriptional changes predominantly occurred in the petals rather than the labellum and staminode.

**Conclusion:** Floral scent evolution in *Cypripedium* is characterized by high chemical diversity, and a weak phylogenetic constraint, consistent with variable selection in food-deceptive systems. The pollination study further suggests the absence of a strong, directed post-pollination effect on the floral scent of *C. henryi*.

## Introduction

Floral volatile organic compounds (VOCs) strongly influence plant reproductive success (Ramya *et al.,* 2020) and play a key role in both short-and long-distance pollinator attraction (Knudsen 2002; Dötterl and Jürgens 2005; B. Song *et al.,* 2015; Dötterl and Gershenzon 2023). More than 1700 floral VOCs have been identified to this date, with Orchidaceae being the most extensively studied plant family (Knudsen *et al.,* 2006; Dötterl and Gershenzon 2023). Through specialized olfactory receptors, insects can interpret floral volatiles as informative chemical cues, guiding them toward specific resources ranging from nutritive rewards like nectar and pollen, to non-nutritive benefits such as brood sites, fragrance compounds or shelter. Over time, foraging insects can learn to associate specific volatile profiles with the presence of food rewards, adapting their foraging behaviour to maximize energetic gain (Haverkamp *et al.,* 2018). Some plants have evolved to exploit these established sensory biases and attract visitors through deceit, using olfactory and/or visual cues without offering the expected reward most often associated with these signals. Orchidaceae contains more deceptive taxa than any other plant family, with roughly one-third of its species relying on some form of deceit, predominantly generalized food deception followed by sexual deception, to achieve pollination (Jersáková *et al.,* 2006). A prominent example is the subfamily Cypripedioideae, commonly known as “slipper orchids”. Species of this orchid lineage are characterized by a distinctive slipper-shaped labellum, which functions as a one-way trap for pollinators. To exit the flower, trapped insects must climb out through the back of the labellum, thereby touching first the stigma and subsequently the pollinia. When the visitors are of appropriate size and shape, and carry a pollen smear on their thorax, this process results in effective pollination (Cribb 1997).

*Cypripedium* L. is a monophyletic genus within Cypripedioideae (J. Li *et al.,* 2011) comprising 54 recognized species (World Checklist of Selected Plant Families, accessed June 2026). This genus is a compelling system for studying scent evolution in deceptive plants as its high diversity of floral VOCs is complemented by increasingly clarified phylogenetic relationships. Cribb and Frosch (2012) recognized 13 sections within *Cypripedium* (Figure 1), all of which have been shown to be monophyletic in phylogenetic studies employing nuclear and chloroplast markers (Fatihah *et al.,* 2011; J. Li *et al.,* 2011). The monophyly of these sections, as well as the topological position of most of the sections, was further supported by incorporating plastome data into a more extensive nuclear loci dataset (41 loci) (Liao *et al.,* 2024). Lagou *et al*. (2024) applied a phylogenomic approach using 913 nuclear loci to resolve remaining uncertainties in the phylogeny of *Cypripedium*. Their results confirmed earlier findings, including sectional monophyly, the position of sect. *Irapeana* as the earliest-diverging lineage, and incongruence between the two traditionally recognized subsections within sect. *Cypripedium* and the two clades actually recovered. Despite the expanded dataset, support along the backbone of the phylogeny remained low, which was attributed to incomplete lineage sorting associated with rapid radiation events, and possible reticulation (Lagou *et al.,* 2024).

**Figure 1.**
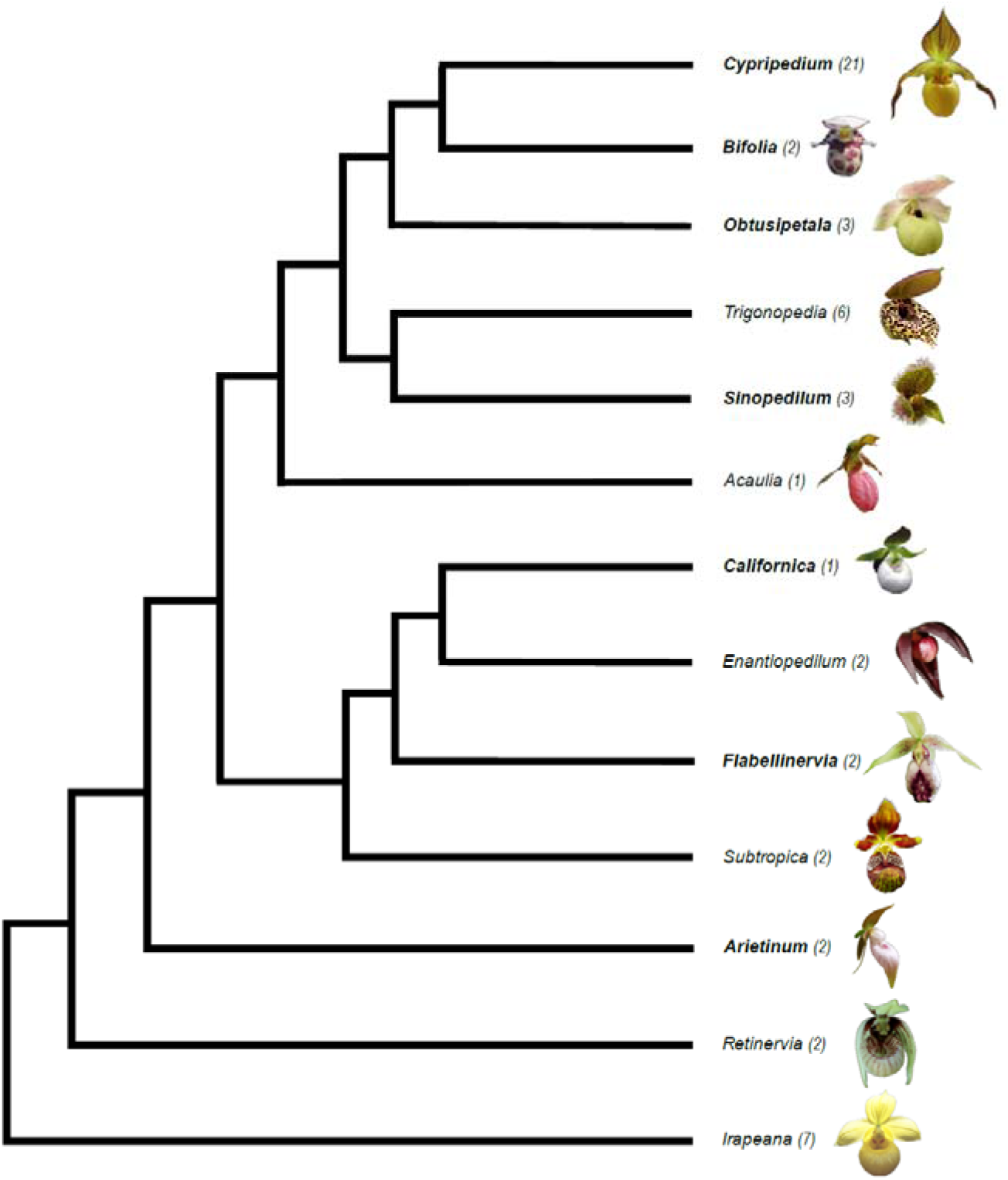
Section-level Cladogram of the genus *Cypripedium*. The cladogram is adapted from the phylogenetic analysis of Lagou et al. (2024), with sectional classification following Frosch and Cribb (2012). Numbers in parenthesis indicate the total number of species currently recognized within each section according to Plants of the World Online (POWO). Sections covered in this study are highlighted in bold. Representative flower morphology for each section is illustrated to the right of the section name: *C. parviflorum* var. *pubescens* f. *planipetalum* (sect. *Cypripedium*), *C. guttatum* (sect. *Bifolia*), *C. flavum* (sect. *Obtusipetala*), *C. lentiginosum* (sect. *Trigonopedia*), *C. micranthum* (sect. *Sinopedilum*), *C. acaule* (sect. *Acaulia*), *C. californicum* (sect. *Californica*), *C. palangshanense* (sect. *Enantiopedilum*), *C. japonicum* (sect. *Flabellinervia*), *C. subtropicum* (sect. *Subtropica*), *C. plectrochilum* (sect. *Arietinum*), *C. debile* (sect. *Retinervia*), *C. irapeanum* (sect. *Irapeana*). Photo credits (from top to bottom): Forest Service of the Northern Region USA, L. Chen, J-B. Chazalon, S. Urban, V. Steindl, V. Steindl, S. Urban, S. Urban, L. Chen, W. Frosch, J-B. Chazalon, S. Urban, M. Béhar.

Research on the genus *Cypripedium* has resulted in documented pollinator records for more than half of the known species. Most species within the genus are pollinated by solitary bees, which likely visit the flowers in search of edible rewards. A notable exception is *C. fasciculatum* Kellogg (sect. *Enantiopedilum*), which is pollinated exclusively by *Cinetus sp.* wasps (Ferguson and Donham 1999). These parasitic wasps are likely attracted to the flower while searching for their fungus gnat hosts. Beyond Hymenoptera, dipteran pollination represents a significant evolutionary strategy within the genus. Sections *Trigonopedia* and *Sinopedilum* have transitioned exclusively to myophily, exhibiting specialized morphological and olfactory traits. These include mottled, dull-coloured foliage and floral volatiles that mimic fermentation or decaying organic matter to lure specific fly families. Recorded pollinators include hoverflies (Syrphidae), fruit flies (Drosophilidae), dung flies (Scathophagidae), blowflies (Calliphoridae), and flat-footed flies (Platypezidae). Dipteran pollination has also been documented in sect. *Obtusipetala* (Vogt 1990; Bänziger *et al.,* 2008; Zheng *et al.,* 2011), sect. *Subtropica* (Jiang *et al.,* 2020), *C. calceolus* of sect. *Cypripedium* (Braunschmid *et al.,* 2017), and has been proposed for *C. debile* of sect. *Retinervia* (Stoutamire 1967; Cribb 1997). Pollinator assemblages may vary geographically: species with broad distributions, such as *C. calceolus* and *C. reginae*, exhibit a greater diversity of pollinators. For example, *C. reginae* has been observed being pollinated by bees in Ontario (Canada) and Missouri (USA), but by hoverflies in Vermont (USA) (Guignard 1886; Vogt 1990; Edens-Meier *et al.,* 2011).

The genus *Cypripedium* is thought to rely primarily on generalized food deception as its pollination strategy (Nilsson 1979; Pemberton 2013), displaying general food related cues to attract naive foragers rather than mimicking a specific model species. However, alternative deceptive mechanisms have been hypothesized for certain species of the genus. For instance, *C. macranthos* var. *rebunense* Kudô may exhibit floral mimicry of *Pedicularis schistostegia* Vved., potentially exploiting pollinators that typically visit that nectar-rewarding species (Sugiura 2002). Similarly, *C. tibeticum* King ex Rolfe is believed to mimic nesting sites of *Bombus* queens, in addition to employing food deception as an attractant mechanism to target the same visitors (P. Li *et al.,* 2006). A rewarding mimicry system was previously described for *C. subtropicum* where hairy tufts on the labellum did not only act as visual lures but also served as food rewards for visiting hoverflies, with a nutrient content comparable to the honeydew secreted by aphids (Jiang *et al.,* 2020). In recent years, guild mimicry has been proposed in Orchidaceae as an intermediate pollination system between generalized food deception and Batesian floral mimicry, where deceptive species do not imitate flowers of a single specific species but rather a group of often distantly related co-flowering species (Jersáková *et al.,* 2016; Scaccabarozzi *et al.,* 2018). Etl *et al*. (2025) proposed that *C. calceolus* employs a mixed pollination strategy, in which guild mimicry plays a significant role. In addition to producing common floral volatiles and bee pheromones, the species emitted more specialized floral compounds also present in the fragrances of multiple co-flowering species.

Although floral scent diversity in Cypripedioideae remains only moderately explored, *Cypripedium* as a genus is comparatively well studied. Early work focused on the European species *C. calceolus* and several North American taxa (Nilsson 1979; Bergstrom *et al.,* 1992; Barkman *et al.,* 1997). Nilsson (1979) proposed that an acetate-rich floral fragrance, comprising compounds such as octyl, decyl, and hexyl acetate, mimics bee volatiles to function as a landing-control pheromone. Bergström *et al*. (1992) confirmed the high acetate proportion in the floral scent of *C. calceolus* but also showcased very different floral fragrance composition of the very closely related *C. parviflorum* var. *parviflorum*, and *C. parviflorum* var. *pubescens*, with different terpenoids together with 1,4-dimethoxybenzene dominating the former and 1,3,5-trimethoxybenzene dominating the latter. Studies on Chinese *Cypripedium* species further illustrate high chemical diversity. In *C. tibeticum*, ethyl acetate, perceived as a faint fruity scent, was suggested to attract bees via food deception (P. Li *et al.,* 2006). The fragrance of *C. plectrochilum* is dominated by 3-methyl-decen-2-one (P. Li *et al.,* 2008a), while *C. henryi* emits mainly fatty acid derivatives, such as dodecanal, undecane, nonanal, and decanal (P. Li *et al.,* 2008b). More recent work by Braunschmid *et al*. (2017, 2020) analysed the scent of *C. calceolus* very thoroughly identifying over 80 floral compounds, making it the species with the best characterized floral scent in the genus. Collectively, available studies highlight pronounced interspecific variation in floral scent composition within *Cypripedium*.

While the effect of pollination on different variables, such as flower life span (Proctor and Harder 1995), colour change (Weiss 1995) and scent emission (Schiestl and Ayasse 2001), have been studied, little is known about the link between pollination-induced changes in VOC composition and the biosynthetic and regulatory pathways involved in scent production and emission. One way to address this question is to examine the transcriptional response of plants by assessing differential gene expression under various pollination treatments. This allows for a better understanding of the molecular basis of how plants mediate their relationships with pollinators and helps to elucidate the processes involved in the evolution of deceptive pollination. For example, the floral scent of sexually deceptive *Ophrys sphegodes* (Orchidaceae) chemically simulates the sex pheromones of its specific pollinator, with alkenes constituting important compounds shared by both the pheromones and the floral scent. It was found that expression of a gene involved in alkene biosynthesis increases in association with increased alkene production (Schlüter *et al.,* 2011). Mutation in such a gene may result in a change in scent composition and thus alter pollinator attraction. Since pollination in *Ophrys* is highly species-specific, this could lead to reproductive isolation and, finally, speciation.

The extensive floral scent diversity within the genus *Cypripedium* offers a unique opportunity to study trait evolution, yet the gene expression profiles that regulate these signals during the floral lifespan remain largely uncharacterized. By characterizing this broad chemical diversity alongside the gene expression patterns triggered by pollination, we can better understand how both historical phylogeny and immediate physiological responses shape the deceptive signals of this genus. Additionally, combining the comprehensive volatile data collected in this study with available species-level phylogenies allows us to evaluate the extent of phylogenetic signal in floral scent phenotypes, which we expect to evolve rapidly in a general food-deceptive lineage such as *Cypripedium.* Here, we generate data of floral VOCs for 31 *Cypripedium* taxa and use these data together with the available phylogeny (Lagou *et al.,* 2024) and pollinator data of the genus to answer the following research questions: (1) How diverse are the floral VOC profiles across the genus *Cypripedium*? (2) How extensive is interspecific variation in floral scent across the genus, and what insights can be gained into its intraspecific variation? (3) Does interspecific scent variation follow a predictable phylogenetic signal, or does it align more closely with shifts in pollinator groups? (4) Finally, does pollination trigger tissue-specific changes in floral volatile production and gene expression within the genus *Cypripedium*? To address this final question, we analysed post-pollination shifts in VOC composition in *C. henryi* across four treatments and linked these chemical changes to underlying gene expression patterns using RNA-seq data from distinct floral tissues.

## Material and Methods

### Plant material

All plant material utilized in this research originated from the living collection of *Cypripedium* maintained and curated at the Oberhof Garden, Eurasburg, a facility associated with the Botanical Garden Munich-Nymphenburg, Germany. A complete list of all studied taxa, with details regarding their specific provenance, is provided in Supplementary Data Table S1. Plants were cultivated in pots either in a greenhouse or outdoors depending on the species requirements. For the genus-wide comparative study, a total of 31 *Cypripedium* taxa, including 24 different species, three natural hybrids and three varieties each of *C. macranthos* and *C. parviflorum*, were included. For the pollination study, two multiflowered clonal individuals of *C. henryi* were sampled (Accession No. 2023/1153).

### Literature review on pollinator species of the genus *Cypripedium*

To establish a comprehensive dataset of *Cypripedium* pollinators, we updated the checklist provided by Pemberton (2013). Our revised dataset includes pollinator entries for six additional *Cypripedium* species previously undocumented in the 2013 review (Supplementary Data Table S2). The review was carried out primarily using Google Scholar, complemented by searches on ResearchGate and JSTOR, and was completed in December 2025. Literature searches were conducted using the keywords “*Cypripedium* pollination” and “*Cypripedium* pollinators.”

### Scent collection

Scent collection for the comparative study was conducted *in situ* at the Oberhof Garden workspace outside the greenhouses. Plant volatiles were collected using a dynamic headspace method following Dötterl *et al*. (2005). Prior to sampling for comparative analyses across the genus, flowers were examined to confirm the presence and intactness of both pollinia. Leaf samples serving as controls were collected for each floral sample (from the same individual) and ambient air samples were also taken on each collection day. Single flowers and leaves were enclosed in polyester oven bags (Toppits R©), and following a scent accumulation of 5 min, samples were collected for 5 min at a flow rate of 0.15 - 0.2 l/min into thermal desorption tubes (TD-tubes) using a pump (Rotary Vane Pump Series G1, Thomas R©). Subsequently, a second sample was collected from the same flower and leaf using the same protocol, but with an extended accumulation and collection time of 15 min each. These two different accumulation and collection times were used as the amounts of scents released from the flowers were unknown and to increase the likelihood of obtaining samples that contain scent amounts well suited for GC/Ms-analyses (see below). Scent collection for the comparative study was carried out between April 27th and May 23^rd^, 2022, between 9 am and 3 pm. The average ambient temperature during scent collection was 18.6 ± 3.1 °C (mean ± SD), with temperatures ranging from 14 to 24 °C. The age of the flowers from which the scent was collected ranged from two to nine days. The flowers of *Cypripedium* last for between two to five weeks (Cribb 1997).

To test for post-pollination effects of scent in *C. henryi*, scent was collected from May 7th to May 16th, 2023, between 10 am and 2 pm under controlled conditions in a greenhouse at the Munich-Nymphenburg Botanical Garden. Here, accumulation and collection time were set at 6 minutes each, based on data obtained in the previous analyses. Polyester oven bags of identical dimensions (15×15 cm) were used for bagging to ensure consistency across all samples. The opening of the flower buds was monitored and the first day the flower opened was set as day zero. A flower was considered as open when the lateral petals, the top sepal and the synsepalum had spread away from the labellum and reached their final position. Plant volatiles of both flower and leaf were collected on day two for the first time. Pollination treatments (see below for details) were carried out directly after the first scent collection. A second scent collection was performed on day four, i.e., two days after the pollination treatments. The average ambient temperature during the first scent collection was 22.2 ± 1.6 °C (mean ± SD), with temperatures ranging from 19 to 25 °C, whereas during the second scent collection it was 21.1 ± 2.0 °C (mean +-SD), ranging from 19 to 24 °C.

### Scent analysis

Scent samples were analysed through Gas Chromatography-Mass Spectrometry (GC/MS). An automatic thermal desorption (TD) system (TD-20, Shimadzu, Japan) was used, coupled to a Shimadzu GC/MS-QP2010 Ultra equipped with a ZB-5 fused silica column (5% phenyl polysiloxane; 60 m, i.d. 0.25 mm, film thickness 0.25 μm, Phenomenex). When loading the samples onto the column, split ratios varied between 1:1 to 1:25 (Supplementary Data Table S1), with a split ratio of 1:1 being used for the *C. henryi* samples of the pollination study. Mass spectra were taken at 70 eV (EI mode) from 34 to 350 m/z. To process the GC/MS data, the GCMSolution package Version 4.50 was used (Shimadzu Corporation 1999-2018). For additional GC/MS setting details see Braunschmid *et al*. (2017). In the comparative study, samples with the shorter accumulation and collection time (5 min) were analysed first. Samples with the longer accumulation and collection time (15 min) were only analysed if the sample with shorter collection time showed a low amount of fragrance compounds. If both samples were analysed, the one with the higher scent amounts (typically the sample with the longer collection time) was selected for downstream analyses. Plant volatiles were identified by database comparisons (retention index, mass spectra), using the databases ADAMS, ESSENTIALOILS-23P, FFNSC 2, W9N11, and a database created by the Plant Ecology Laboratory of the University of Salzburg using available synthetic standards. A compound was identified as a floral volatile if it was absent in the leaf sample, or if the peak area (total ion current) in the gas chromatogram was at least five times higher in the flower sample than in the associated leaf sample. In the experimental pollination study with *C. henryi*, floral compounds with a single occurrence were excluded from the analyses.

### Pollination treatments of *Cypripedium henryi* flowers

Four flower-level treatments were conducted using 12 flowers from two plant individuals. (1) Removal of both pollinia (removal); (2) deposition of a pollinium from the same clonal accession on the stigma (deposition); (3) both the removal and deposition treatment combined (deposition+removal); (4) control group with no artificial pollination performed (control). To minimize potential plant-individual bias, flowers assigned to each treatment were distributed as evenly as possible across the two sampled plants. Due to a delayed onset of volatile emission, two flowers (one each from the control and removal group) were excluded from the scent analysis. Consequently, these samples were also omitted from the differential expression analysis and all subsequent steps within the transcriptomic pipeline.

### RNA isolation and Transcriptome Sequencing

The labellum, staminode and lateral petals were collected in liquid nitrogen immediately following the second scent sampling (i.e. 48 h (± 2 h) after pollination treatment). Total RNA was isolated using the InnuPREP Plant RNA Kit (Innuscreen GmbH). For each flower, the whole staminode (around 15 mg), one longitudinal half of the labellum (around 60 mg) and a whole lateral petal (around 30 mg) were used for RNA extraction. Prior to the extraction, the tissue samples were crushed on dry ice using a rotor-stator homogenizer. Total RNA was eluted in 20 µl RNAse-free H_2_O. RNA quality was assessed via Agilent ScreenTape, with the RIN threshold of samples ranging between 5.8 and 8.4. Both transcriptome library preparation and sequencing were performed by BMKGENE (Münster, Germany). Libraries were constructed using a poly(A)-selection strategy and a strand-specific (dUTP) protocol, followed by sequencing on an Illumina NovaSeq 6000 platform to generate 150 bp paired-end reads.

### Transcriptome assembly

Pre-assembly of transcriptomic data was performed following the workflow of Siadjeu and Kadereit (2024). The following tools were used: FastQC v0.12.1 (https://www.bioinformatics.babraham.ac.uk/projects/fastqc/) and MultiQC v1.17 (Ewels *et al.,* 2016) for sequence quality control assessment, Rcorrector v1.0.6 (L. Song and Florea 2015) for correction of sequencing errors, TranscriptomeAssemblyTools (https://github.com/harvardinformatics/TranscriptomeAssemblyTools) for removal of uncorrectable reads, TrimGalore v0.6.10 (https://github.com/FelixKrueger/TrimGalore/releases) for filtering of adapters and low quality reads, and finally Bowtie2 2.5.2 (Langmead and Salzberg 2012) and SILVA v-138 rRNA database for removal of rRNA.

De novo transcriptome assembly was performed using Trinity v2.15.1 (Grabherr *et al.,* 2011) with the following parameters: Trinity –seqType fq –max_memory 150G –CPU 54. The quality of the transcriptome assembly was checked using two different methods. First read support was checked by mapping the cleaned reads to the assembled transcriptome using Bowtie2 v2.5.2. Second BUSCO completeness was checked by using the software BUSCO v5.5.0 (Manni *et al.,* 2021).

### Transcript quantification and differential expression analysis

The assembled reads were clustered at 98% similarity with CD-HIT v4.8.1 (Fu *et al.,* 2012) to remove duplicates and obtain the final transcripts. For subsequent analysis, transcripts with an ORF were extracted using Transdecoder v.5.5.0 (https://github.com/sghignone/TransDecoder). Transcript quantification was performed using Salmon 1.10.2 (Patro *et al.,* 2017). Only transcripts with an occurrence of at least ten in three different samples were kept for further analysis. Differential expression analysis was performed in R Statistical Software v4.3.3 (R Core Team 2025) using the DESeq2 package (Love *et al.,* 2014). To account for inter-individual variation between the two sampled plants, Plant ID was incorporated into the generalized linear model as a blocking factor (design = ∼ Plant_ID + Treatment). Transcripts were considered differentially expressed if they exhibited a false discovery rate (FDR) of <5% (adjusted p-value <0.05) and a ∣log_2_FC∣>1. To evaluate the impact of pollination on volatile-related gene expression, each treatment group was compared against the control across three distinct floral tissues: petals, staminodes, and labella. For the purpose of this analysis, “petals” refers exclusively to the lateral petals, as distinct from the labellum, which represents a morphologically modified third petal.

### Gene annotation and candidate gene selection

Metabolic pathways and annotations of differentially expressed transcripts were assigned via the tool Mercator4 v6.0 (Schwacke *et al.,* 2019). Both swiss-prot protein sequences database and prot-scriber were included to improve the annotations. Subsequent analysis focused on genes involved in floral volatile biosynthesis in *C. henryi*, specifically those associated with the fatty acid (FA) and terpenoid biosynthesis pathways.

### Data visualisation and statistical analysis

To visualize scent sample (dis)similarity, NMDS plots were created using the R packages vegan v2.6-4 (Oksanen *et al.,* 2024) and ggplot2 v4.0.1 (Wickham 2016). For the genus-wide dataset, we additionally utilized the package chemodiv v0.2.0 (Petrén *et al.,* 2023) to compute Generalized UniFrac distances, which account for structural and biosynthetic similarity between volatiles. Specifically, Generalized UniFrac (GenUniFrac) dissimilarities were calculated with the parameter α set to 1, thereby weighting compound dissimilarities in direct proportion to their relative abundances. To visualize the distribution of floral VOCs across all sampled *Cypripedium* taxa, a heatmap based on relative abundances was generated using the R package pheatmap (Kolde 2010). To simplify the visualization and focus on major constituents, the dataset for this heatmap was reduced to only include floral VOCs whose total relative quantity across sampled taxa exceeded an arbitrary value of 8%.

Next to determine the proportion of variance in the floral scent composition explained by taxonomic section and pollinator group, we conducted a PERMANOVA (Permutational Multivariate Analysis of Variance) with 10,000 permutations, based on chemically informed GenUniFrac dissimilarities. Dissimilarity matrices were generated using the chemodiv package, and multivariate testing was performed in vegan. We categorized the *Cypripedium* taxa in our dataset into three main pollinator syndrome categories: (1) bees, (2) bees and floricolous flies, (3) drosophilid flies. These assignments were derived from primary literature or, in the absence of empirical records, through phylogenetic inference while also considering the visual phenotype of the flower. To assess the correlation between floral scent composition and phylogenetic relationships, we performed a Mantel test across the 31 sampled taxa. This analysis compared the GenUniFrac dissimilarity matrix against a matrix of pairwise phylogenetic distances, which were derived from the Lagou *et al*. (2024) phylogeny using the ape v5.8 package.

To assess the effect of pollination treatment on the floral scent of *C. henryi* while accounting for individual variation, we conducted a Type-I sequential PERMANOVA. Treatment type was used as a fixed factor, with Plant ID entered first in the model to partition out inter-individual variance. Again, this analysis was based on GenUniFrac dissimilarities calculated with chemodiv. Additionally, paired t-tests were conducted to assess variation within individual flowers between the first and second scent collection events for both the total scent emission and the relative abundances of individual compounds. Since our analysis did not include internal calibration standards, total scent emission was estimated as the sum of all volatile peak areas. Although absolute emission rates were not quantified, total peak area serves as a reliable proxy for total scent output, allowing valid intra-individual comparisons given our standardized sampling conditions. For compound-level analyses, p-values were corrected for multiple testing using the Bonferroni method. Expression profiles of the differentially expressed candidate genes identified across pollination treatments in *C. henryi* were visualized using the R package pheatmap (Kolde & Kolde, 2015).

## Results

### Floral fragrance composition in *Cypripedium*

In the 44 analysed scent samples collected from 31 *Cypripedium* taxa, a total of 258 floral volatiles were found of which 174 were (tentatively) identified. These included 64 aliphatic compounds, 63 terpenoids, 34 aromatic compounds, eight C5-branched compounds, three N-bearing compounds, as well as one miscellaneous cyclic compound and one spiroacetal (Supplementary Data Table S3). The total number of floral compounds detected per flower ranged from 1 to 42 compounds, with an average of 14.1 ± 12.7 (SD) per sample and 15.2 ± 12.8 (SD) at the taxon level. Floral scent samples of *C. calceolus* and *C. californicum* held the highest number of floral volatiles with 42 and 39 compounds, respectively. The floral volatile with the highest occurrence was the monoterpenoid linalool (Figure 2), which was identified as a floral compound in nine taxa of sect. *Cypripedium* and in *C. californicum* from the monotypic sect. *Californica*. Linalool derivatives like linalool oxides (furanoids, pyranoids) and lilac compounds, were also detected in smaller amounts than linalool, and only occurred in samples that also included linalool with the exception being *C. froschii*. The other most often occurring floral VOCs were the aliphatic compounds (*Z*)-3-hexen-1-ol (8 taxa) and 1-hexanol (7 taxa), the aromatic compounds benzyl alcohol (7 taxa) and 2-phenylethanol (6 taxa) and the monoterpenoid (*E*)-β-ocimene (7 taxa). Identification was limited (≤75% of total scent) for five of the 31 taxa studied. Most notably, *C. flavum* exhibited a particularly complex profile of unknowns, with 20 compounds remaining unidentified (comprising 27% of its total scent). In contrast, the unidentified portion of the *C. passerinum* bouquet (29%) consisted of only two components. Two further unidentified compounds (unk_739 and unk_1097) contributed significantly to the profiles of *C. froschii* (9% and 21.4%), *C. macranthos* var. *hotei-atsumorianum* (7.1% and 10.5%), and *C. macranthos* var. *rebunense* (8.1% and 10.5%). Detailed RI and mass spectral data for all unidentified components are available in Supplementary Data Table S4.

**Figure 2.**
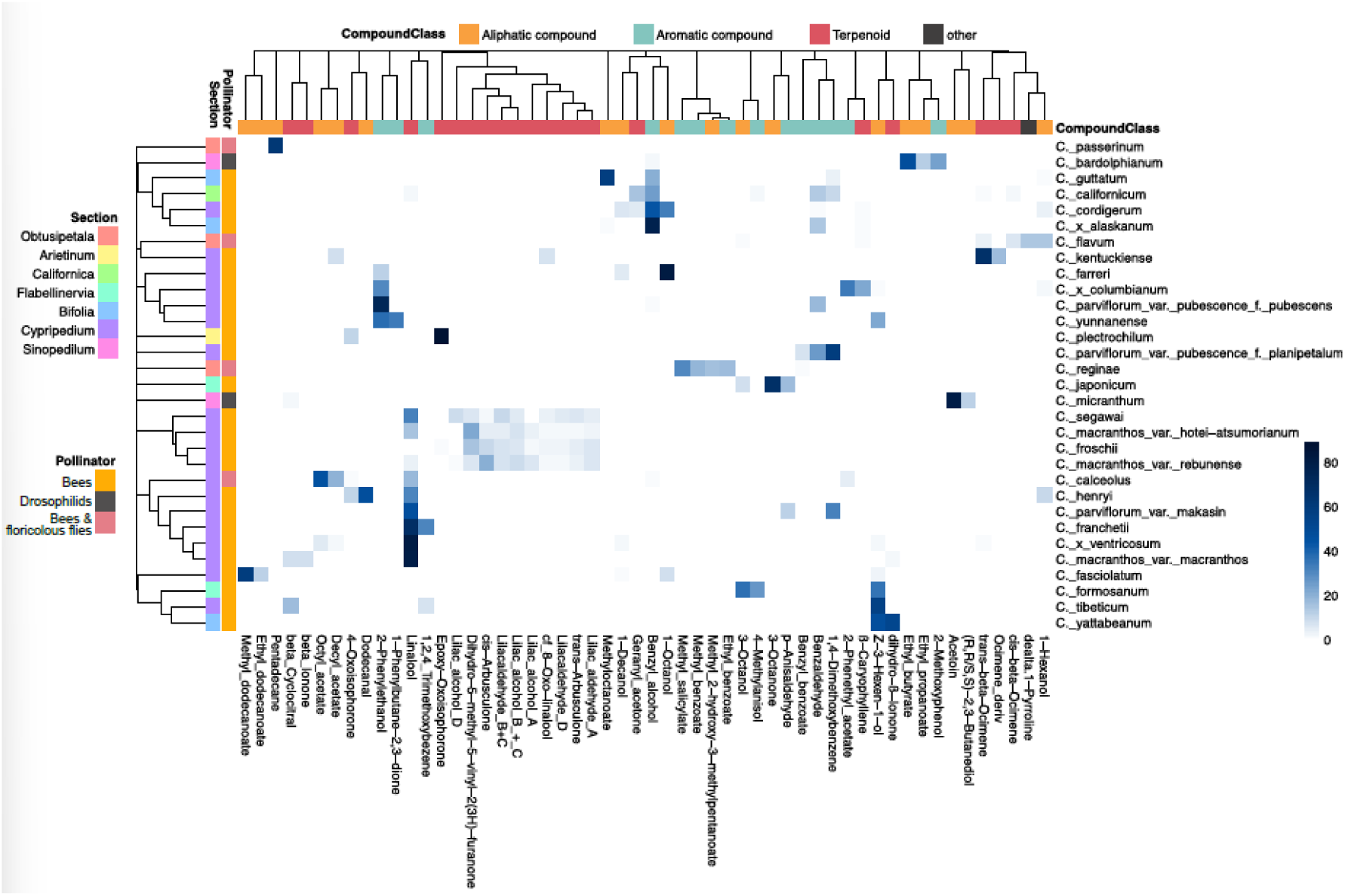
Floral volatile occurrence across sampled *Cypripedium* spp. The blue colour gradient represents the relative proportion of specific floral VOCs within each species scent profile. For better visualization the dataset was reduced to only include floral VOCs whose relative quantity across samples exceeded 8%.

### Comparison of floral scent across *Cypripedium* sections

The average number of detected floral volatiles per taxon (n = 31; comprising 24 species and 6 varieties) as well as the chemical nature of compounds varied notably among the different sections of *Cypripedium* covered in this study (Figure 3). Section *Arietinum* exhibited the lowest floral volatile diversity, with only two compounds, epoxy-oxoisophorone (89%) and 4-oxoisophorone (11%), both terpenoids, detected in the single sampled taxon, *C. plectrochilum*. Section *Bifolia* averaged 10.7 floral volatiles per sampled taxon and consisted of aliphatics, terpenoids and aromatic compounds. Only two floral compounds were identified in *C. yatabeanum*, the terpenoid dihydro-ß-ionone (53%) and the aliphatic compound (*Z*)-3-hexen-1-ol (47%). The floral scents of the other taxa of sect. *Bifolia* were more complex: *C.* x *alaskanum* was dominated by the aromatic compounds benzyl alcohol (78%) and benzaldehyde (15%), with minor aliphatic and terpenoid components, while *C. guttatum* had the most diverse profile, including the aliphatic compound methyl octanoate (58%), the aromatic compounds benzyl alcohol (21%), 1,4-dimethoxybenzene (4%) and methyleugenol (3%), and the terpenoid β-citronellol (2%). The highly complex scent of *C. californicum* of the monotypic sect. *Californica* was composed almost exclusively of aromatics and terpenoids with benzyl alcohol (25%), geranyl acetone (15%) and methoxymethylbenzene (14%) being the most prevalent compounds. Section *Cypripedium*, to which the bulk of the sampled taxa belonged, averaged 8.8 floral volatiles per taxon. Aliphatics, aromatics, and terpenoids were about equally prevalent in this section, alongside a notable proportion of unidentified compounds. Linalool was the most prominent VOC, occurring in nearly half of the section’s taxa (9 out of 19) and acting as the dominant scent constituent in five of those cases. Floral scent of sect. *Flabilinervia* consisted of only six different compounds, with exclusively aliphatics and aromatics making up the scent of the two species of this section. Floral scent of *C. japonicum* was dominated by 3-octanone (68%) and *p*-anisaldehyde (17%) while the scent of *C. formosanum* flowers was more evenly distributed around 3-octanol (37%), (*Z*)-3-hexen-1-ol (36%) and 4-methylanisole (27%). Flowers of sect. *Obtusipetala* exhibited the chemically most diverse scent profile with representatives of most major compound classes, and also notable quantities of C5-branched chain compounds, N-bearing compounds and a significant proportion of unknown volatiles. This diversity was reflected by the dominant compounds of the three sampled taxa: the aliphatic pentadecane (63%) in *C. passerinum*, the aromatic methyl salicylate (31%) in *C. reginae*, and the N-bearing δ-1-pyrroline (15%) in *C. flavum*. Finally, both sampled species of sect. *Sinopedilum*, *C. micranthum* and *C. bardolphianum*, had floral scents dominated by aliphatic compounds, with *C. micranthum* primarily emitting acetoin (80%) and the floral scent of *C. bardolphianum* being characterized by ethyl butyrate (48%) and the aromatic compound 2-methoxyphenol (24%).

**Figure 3.**
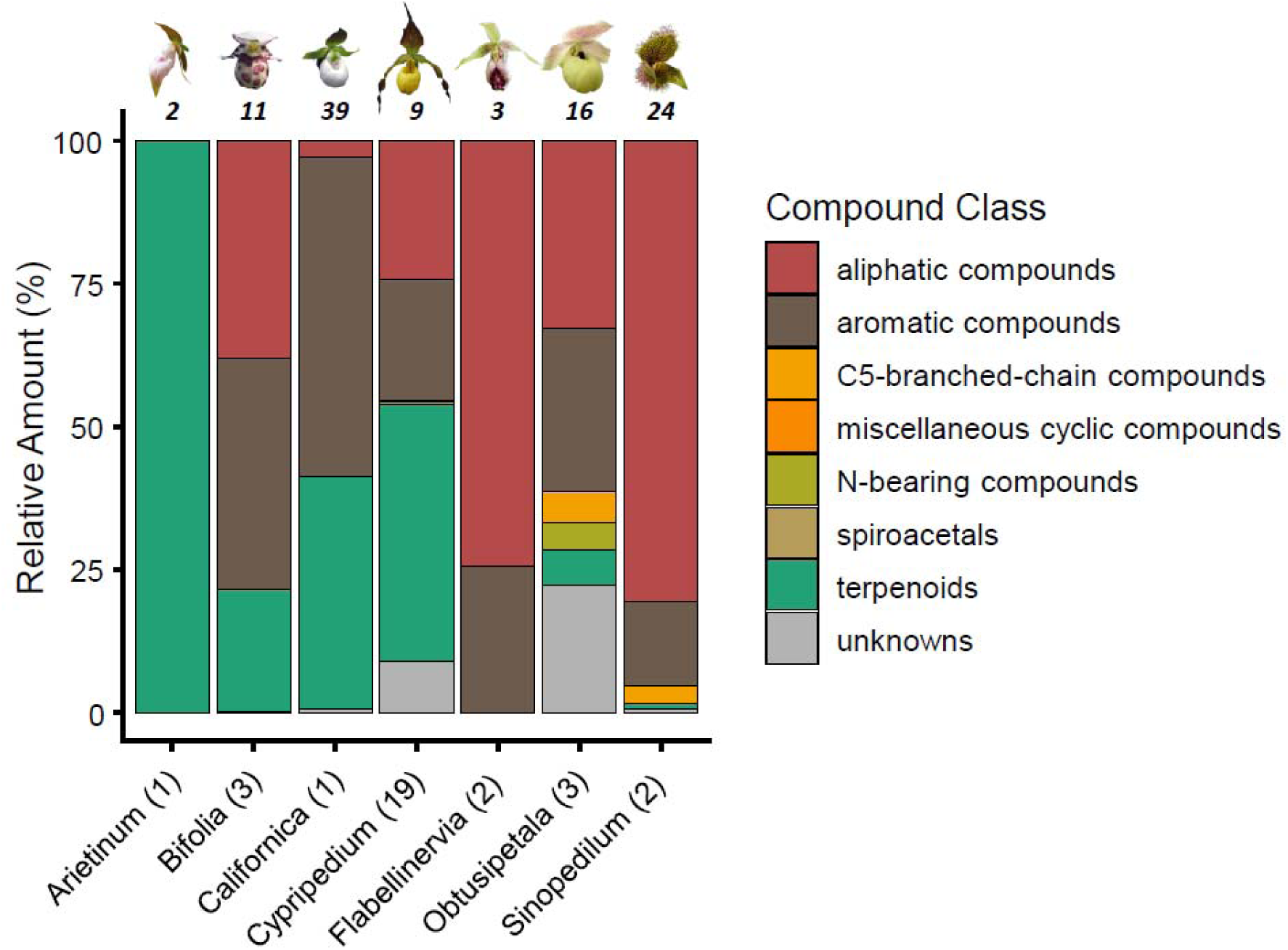
Mean relative abundance (%) of floral volatile compound classes across sampled *Cypripedium* sections. The number of taxa sampled within each section is provided in parentheses. Values above the bars represent the rounded mean number of total floral volatiles identified per taxon for that respective section. Representative flowers for each taxonomic section are shown above the bars: *C. plectrochilum* (sect. *Arietinum*), *C. guttatum* (sect. *Bifolia*), *C. californicum* (sect. *Californica*), *C. parviflorum* var. *pubescens* (sect. *Cypripedium*), *C. japonicum* (sect. *Flabellinervia*), *C. flavum* (sect. *Obtusipetala*) and *C. micranthum* (sect. *Sinopedilum*). See Fig. 1 for credits of the flower images.

### Floral scent variation within *Cypripedium*

Floral scent varied also within each section of *Cypripedium* (Figure 4). We found substantial diversity in floral volatiles among *Cypripedium* species, with many compounds being unique to a single taxon out of the 31 investigated. Notably, acetoin, pentadecane, methyl dodecanoate, and dodecanal constituted respectively 80%, 63%, 59%, and 47% of the total floral fragrance in individual species, while not being identified as floral volatiles in any of the other taxa examined. The PERMANOVA results indicated that the taxonomic sections accounted for 15% of the total scent variation, though this effect was not statistically significant (F_5,23_ = 0.95, p = 0.586). In contrast, pollinator groups significantly explained 10% of the variance in floral scent composition (F_2,23_ = 1.55, p = 0.049). We also detected substantial intraspecific variation of floral scent profiles among the three sampled infraspecific taxa of *C. parviflorum*. Only three floral VOCs were shared across the taxa examined, and none of these occurred in all three. 1,4-Dimethoxybenzene dominated the floral fragrances of both *C. parviflorum* var. *pubescens* f. *planipetalum* (59%) and *C. parviflorum* var. *makasin* (32%). The only shared compound between *C. parviflorum* var. *makasin* and *C. parviflorum* var. *pubescens* f. *pubescens* was indole (5% and 2%, respectively). Finally, benzaldehyde was a major constituent in *C. parviflorum* var. *pubescens* f. *planipetalum* (26%) and *C. parviflorum* var. *pubescens* f. *pubescens* (18%). In comparison, the three sampled taxa of *C. macranthos* exhibited a higher degree of volatile overlap. This was especially the case for *C. macranthos* var *rebunense* and *C. macranthos* var. *hotei-atsumorianum*, which shared a similar overall floral scent profile (Figure 4). The floral scent profile of *C. macranthos* var. *macranthos* consisted of only four VOCs, with linalool being the sole compound shared with the other two sampled infraspecific taxa of the species.

**Figure 4.**
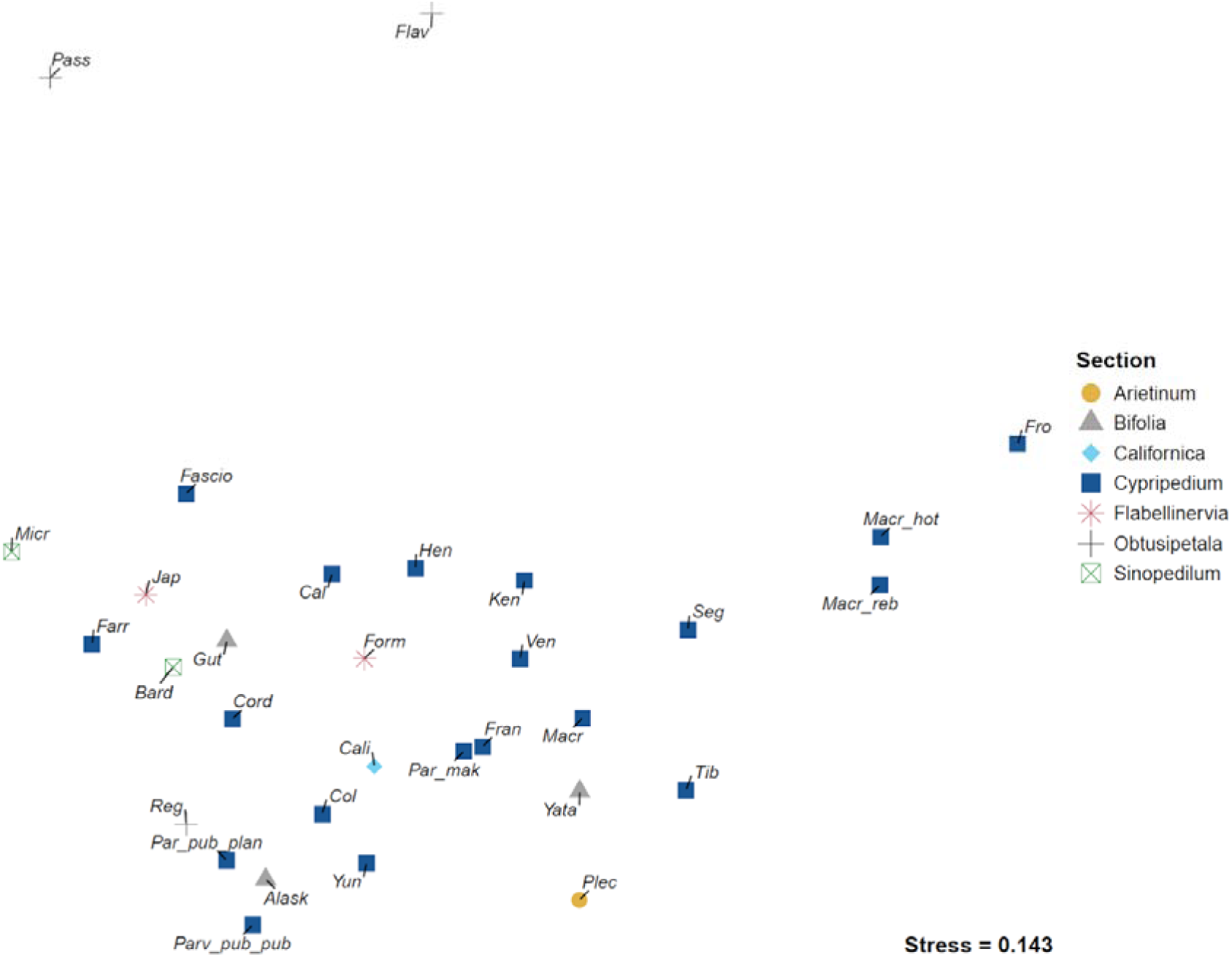
Non-metric multidimensional scaling (NMDS) showing the differences in relative amounts of floral scent components among the 31 sampled *Cypripedium* taxa. Samples are labelled with abbreviations corresponding to their respective taxa.

### Phylogenetic signal analysis in floral scent of *Cypripedium*

A Mantel test revealed a significant positive correlation between scent profile dissimilarity and phylogenetic distance (p = 0.048), indicating that phylogenetically closely related species share more similar scent compositions. However, the effect size measured as the Pearson correlation was low with r = 0.161.

### Post-pollination effect on floral scent of *Cypripedium henryi*

In the 20 analysed floral scent samples (ten each pre-and post-treatment), 28 of the 33 detected floral volatiles could be identified (Table 1). The identified floral volatiles were mostly aliphatics (19 compounds) followed by terpenoids (5 compounds) and aromatic compounds (4 compounds). Five floral volatiles could not be identified but represented a small proportion of less than 1 % of the total floral fragrance. Floral scent samples prior to pollination treatments were dominated by aliphatic compounds, specifically FA derivatives. Dodecanal was the most abundant compound, accounting for 57–77% of the total volatile emission, followed by undecane (4.8–16.8%). Nonanal and decanal were generally the next highest concentrated VOCs in floral scent samples, but were only intermittently identified as floral VOCs, as their abundance did not consistently exceed the five-fold threshold relative to leaf control samples. In addition to these dominant volatiles, a suite of other FA-derived compounds was consistently present in all samples, regardless of treatment. These included hexanoic acid, 1-decanol, and several C12 to C14 derivatives (Table 1). Next to these aliphatic compounds, the terpenes linalool and 4-oxoisophorone were also consistently present, with linalool contributing 2.1–13.3% and 4-oxoisophorone about 1% to the floral scent.

**Table 1.**
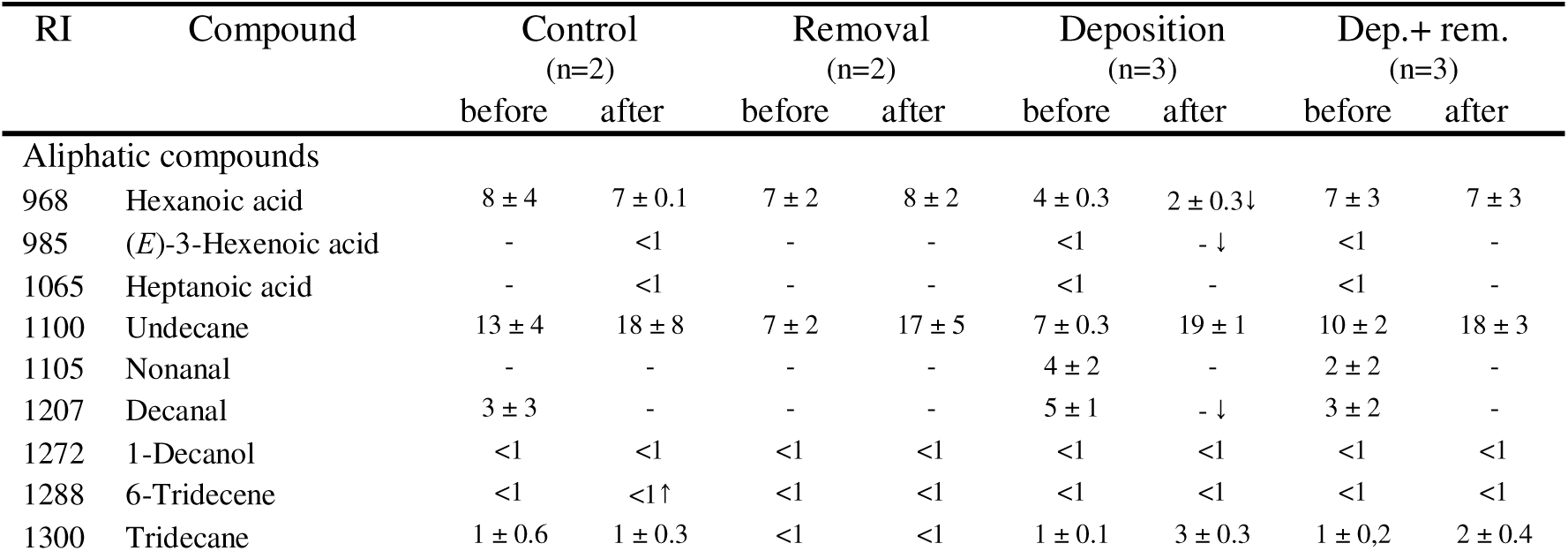
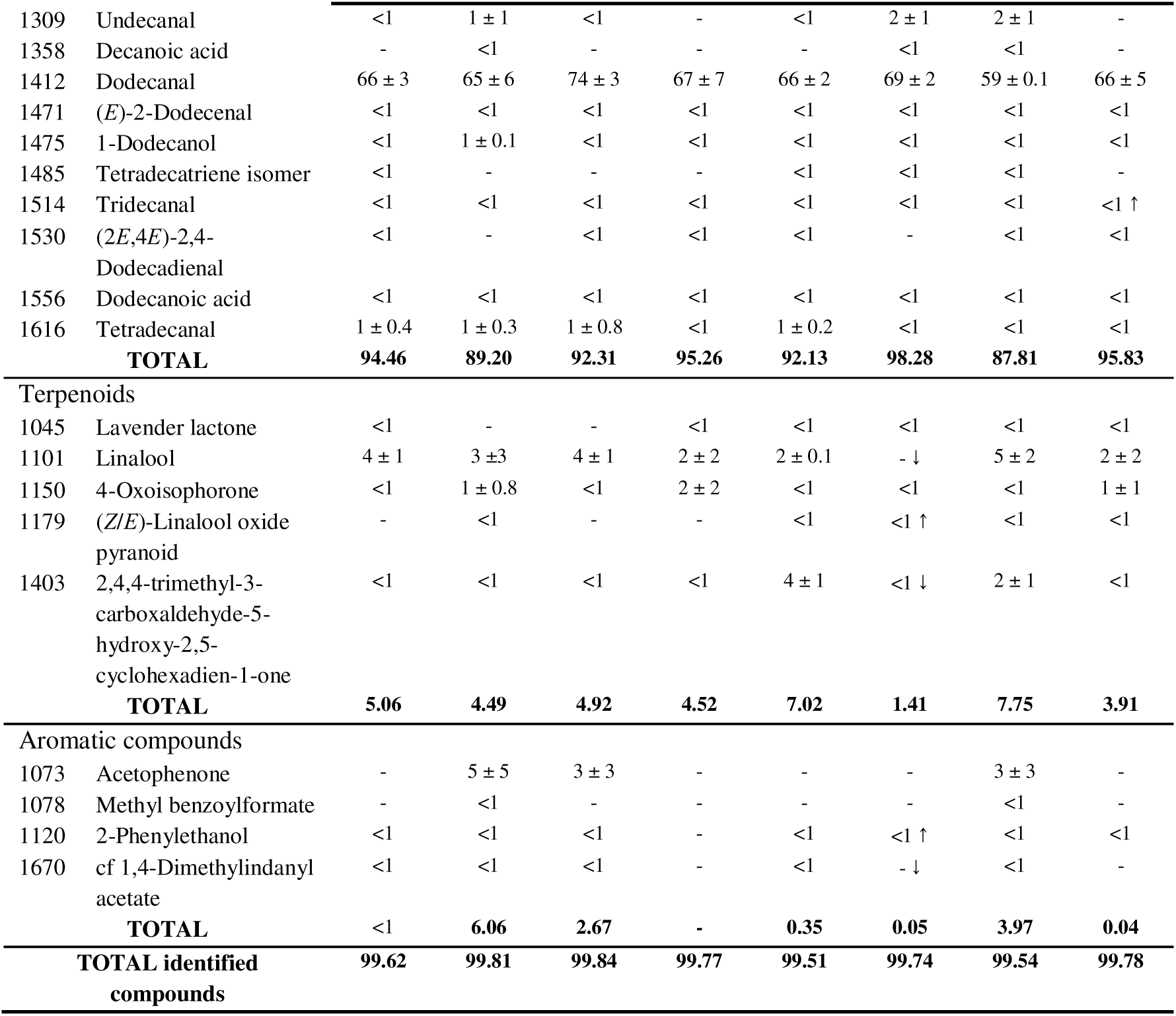
Mean relative amount of *C. henryi* floral volatiles per treatment group before and after pollination treatment application. Control: no pollination; **removal:** pollinia removal; **deposition:** pollination; **dep. + rem.:** pollination and pollinia removal. The standard error is indicated for values higher than one. Arrows indicate statistically significant differences in relative amounts before and after the pollination treatment (paired t-test p < 0.05), prior to the Bonferroni correction.

| RI | Compound | Control |  | Removal |  | Deposition |  | Dep.+ rem. |  |
|---|---|---|---|---|---|---|---|---|---|
|  |  | (n=2) |  | (n=2) |  | (n=3) |  | (n=3) |  |
|  |  | before | after | before | after | before | after | before | after |
| Aliphatic compounds |  |  |  |  |  |  |  |  |  |
| 968 | Hexanoic acid | 8 ± 4 | 7 ± 0.1 | 7 ± 2 | 8 ± 2 | 4 ± 0.3 | 2 ± 0.3↓ | 7 ± 3 | 7 ± 3 |
| 985 | (*E*)-3-Hexenoic acid | - | <1 | - | - | <1 | - ↓ | <1 | - |
| 1065 | Heptanoic acid | - | <1 | - | - | <1 | - | <1 | - |
| 1100 | Undecane | 13 ± 4 | 18 ± 8 | 7 ± 2 | 17 ± 5 | 7 ± 0.3 | 19 ± 1 | 10 ± 2 | 18 ± 3 |
| 1105 | Nonanal | - | - | - | - | 4 ± 2 | - | 2 ± 2 | - |
| 1207 | Decanal | 3 ± 3 | - | - | - | 5 ± 1 | - ↓ | 3 ± 2 | - |
| 1272 | 1-Decanol | <1 | <1 | <1 | <1 | <1 | <1 | <1 | <1 |
| 1288 | 6-Tridecene | <1 | <1↑ | <1 | <1 | <1 | <1 | <1 | <1 |
| 1300 | Tridecane | 1 ± 0.6 | 1 ± 0.3 | <1 | <1 | 1 ± 0.1 | 3 ± 0.3 | 1 ± 0,2 | 2 ± 0.4 |
| 1309 | Undecanal | <1 | 1 ± 1 | <1 | - | <1 | 2 ± 1 | 2 ± 1 | - |
| 1358 | Decanoic acid | - | <1 | - | - | - | <1 | <1 | - |
| 1412 | Dodecanal | 66 ± 3 | 65 ± 6 | 74 ± 3 | 67 ± 7 | 66 ± 2 | 69 ± 2 | 59 ± 0.1 | 66 ± 5 |
| 1471 | (*E*)-2-Dodecenal | <1 | <1 | <1 | <1 | <1 | <1 | <1 | <1 |
| 1475 | 1-Dodecanol | <1 | 1 ± 0.1 | <1 | <1 | <1 | <1 | <1 | <1 |
| 1485 | Tetradecatriene isomer | <1 | - | - | - | <1 | <1 | <1 | - |
| 1514 | Tridecanal | <1 | <1 | <1 | <1 | <1 | <1 | <1 | <1 ↑ |
| 1530 | (2*E*,4*E*)-2,4-Dodecadienal | <1 | - | <1 | <1 | <1 | - | <1 | <1 |
| 1556 | Dodecanoic acid | <1 | <1 | <1 | <1 | <1 | <1 | <1 | <1 |
| 1616 | Tetradecanal | 1 ± 0.4 | 1 ± 0.3 | 1 ± 0.8 | <1 | 1 ± 0.2 | <1 | <1 | <1 |
|  | **TOTAL** | **94.46** | **89.20** | **92.31** | **95.26** | **92.13** | **98.28** | **87.81** | **95.83** |
| Terpenoids |  |  |  |  |  |  |  |  |  |
| 1045 | Lavender lactone | <1 | - | - | <1 | <1 | <1 | <1 | <1 |
| 1101 | Linalool | 4 ± 1 | 3 ±3 | 4 ± 1 | 2 ± 2 | 2 ± 0.1 | - ↓ | 5 ± 2 | 2 ± 2 |
| 1150 | 4-Oxoisophorone | <1 | 1 ± 0.8 | <1 | 2 ± 2 | <1 | <1 | <1 | 1 ± 1 |
| 1179 | (*Z*/*E*)-Linalool oxide pyranoid | - | <1 | - | - | <1 | <1 ↑ | <1 | <1 |
| 1403 | 2,4,4-trimethyl-3-carboxaldehyde-5-hydroxy-2,5-cyclohexadien-1-one | <1 | <1 | <1 | <1 | 4 ± 1 | <1 ↓ | 2 ± 1 | <1 |
|  | **TOTAL** | **5.06** | **4.49** | **4.92** | **4.52** | **7.02** | **1.41** | **7.75** | **3.91** |
| Aromatic compounds |  |  |  |  |  |  |  |  |  |
| 1073 | Acetophenone | - | 5 ± 5 | 3 ± 3 | - | - | - | 3 ± 3 | - |
| 1078 | Methyl benzoylformate | - | <1 | - | - | - | - | <1 | - |
| 1120 | 2-Phenylethanol | <1 | <1 | <1 | - | <1 | <1 ↑ | <1 | <1 |
| 1670 | cf 1,4-Dimethylindanyl acetate | <1 | <1 | <1 | - | <1 | - ↓ | <1 | - |
|  | **TOTAL** | <1 | **6.06** | **2.67** | **-** | **0.35** | **0.05** | **3.97** | **0.04** |
| **TOTAL identified compounds** |  | **99.62** | **99.81** | **99.84** | **99.77** | **99.51** | **99.74** | **99.54** | **99.78** |

Changes in floral scent emission following treatment were predominantly quantitative rather than qualitative, as the chemical composition remained largely consistent across all samples. Total scent generally decreased between collection events; while the groups control, removal, and deposition+removal showed non-significant reductions of 1.9-fold, 2.9-fold, and 3.6-fold, respectively (p > 0.05), the deposition treatment exhibited a significant 3.3-fold reduction (p = 0.003). The decline in absolute amounts was consistent for all major volatiles across all treatment groups. At the individual compound level, several relative shifts were observed within the different treatment groups (see Table 1). However, none of these relative differences remained significant after Bonferroni correction for multiple comparisons. Although not statistically significant, linalool showed a downward trend across all groups, being ubiquitous in the first scent collection but only present in three post-treatment samples (one replicate each from control, removal, and deposition+removal).

NMDS ordination of *C. henryi* floral scent samples showed substantial overlap between post-pollination treatment groups (Figure 5). While samples from the deposition treatment formed a loose grouping post-pollination, a similar pattern was also evident in the corresponding pre-treatment samples. Type-I sequential PERMANOVA of post-treatment scent samples revealed that the pollination treatments had no significant effect on the relative floral scent composition (F_3,5_ = 1.36, p = 0.282). In contrast a highly significant ‘Plant ID’ effect was detected (F_1,5_ = 5.97, p = 0.006), which accounted for 39.7% of the observed variation in scent profiles.

**Figure 5.**
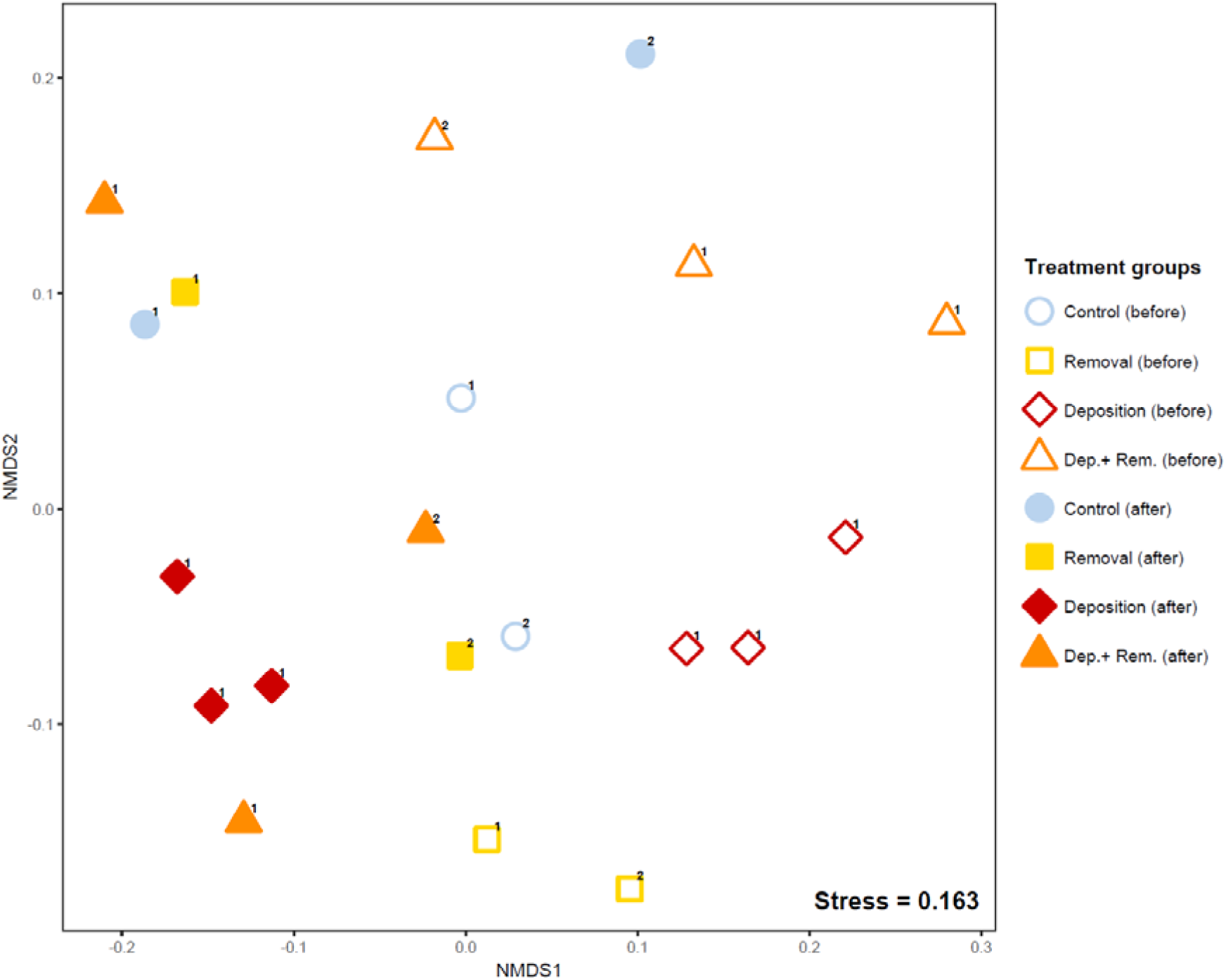
**NMDS analysis visualizing the semi-quantitative floral scent data obtained from floral scent samples of *C. henryi* before and after pollination treatments. Control**: no pollination treatment applied; **Removal:** pollinia removal; **Deposition:** pollination; **Dep.+ Rem.:** pollination and pollinia removal. Open symbols indicate samples collected prior to treatment, whereas filled symbols represent post-treatment samples. Numbers indicate the plant ID of the two sampled plants.

### Transcriptome analysis

RNA sequencing yielded between around 31M and 43M reads per sample. In total, 2,647,661,174 reads were processed with Rcorrector, resulting in the correction of 987,472,741 bases. The de novo transcriptome assembly generated a total of 563,487 transcripts. This study was conducted using de a novo transcriptome assembly that did not have reliable gene models. As a result, we performed expression quantification and differential expression analysis at the transcript level instead of the gene level. Therefore, throughout the manuscript, we will refer to significantly differentially expressed features as differentially expressed transcripts (DETs). Evaluation with BUSCO (n = 1,614, *embryophyta_odb10*) indicated high completeness (C: 95.3%), with 19.6% single-copy and 75.7% duplicated BUSCOs, alongside low fragmentation (2.9%) and few missing genes (1.8%). Read support analysis further confirmed the robustness of the assembly, with 98.91% of reads aligning back to the transcriptome. Using TransDecoder, 124,929 coding transcripts were identified. After filtering for low-abundance transcripts and excluding the samples from the two flowers with delayed volatile emission (one each from the control group and the removal treatment), the dataset used for differential expression analysis consisted of 70,804 transcripts.

### Differential expression analysis

To identify treatment-induced transcriptional shifts, the transcriptomes from treated flowers were compared against the control group within each tissue type. Overall, the petals exhibited a markedly higher number of DETs than the labellum or staminode (Supplementary Table S5). Functional annotation with Mercator4 revealed 19 DETs linked to FA biosynthesis, and a single DET associated with terpenoid synthesis. Consistent with the global expression trends, the transcriptional response of identified candidate genes was more extensive in the petals than in the labellum or staminode (Figure 6). Expression profiles revealed almost no patterns restricted exclusively to the two treatments involving pollination (deposition and deposition+removal). The sole exception was the chloroplastic pyruvate dehydrogenase E1 component subunit alpha-3 (*PDH-E1-*α), involved in acetyl-CoA generation, which was slightly but significantly up-regulated in petals. Furthermore, treatment-specific patterns shared across tissues were rare, with the exception of 3-ketoacyl-CoA synthase 20 (*KCS20*). This gene, involved in FA elongation, was down-regulated following the deposition treatment in both the staminode and labellum, but remained stable in the petals. Several DETs associated with FA biosynthesis, exhibited uniform tissue-specific responses across all treatments. For instance, the up-regulation of *ZC3H55* (encoding zinc finger CCCH domain-containing protein 55) was restricted to the staminode. In contrast, petal-specific up-regulation was observed for two acyl-coenzyme A oxidase paralogs (*ACX3* and *ACX1_1*) and an isoform of biotin carboxylase 1 (*BCCP1_2*), while the peroxisomal long-chain acyl-CoA synthetase 6 (*LACS6*) was consistently down-regulated in the petals across all treatments. Regulation of *BCCP1* was found to be isoform-specific within the petals, where the two identified isoforms showed opposing directions of differential expression. The only identified terpenoid-related DET was 3-hydroxy-3-methylglutaryl-CoA reductase 3 (*HMGCR3*), a key enzyme in the mevalonate (MVA) pathway. *HMGCR3* exhibited a highly tissue-specific response profile: while it was constitutively up-regulated across all treatments in the petals, its activation in the staminode was restricted to both the removal and deposition group. Notably, in the labellum, *HMGCR3* was uniquely up-regulated only under the combined deposition+removal treatment. Detailed statistical values, including log_2_FC and p_adj_, for these transcripts are provided in Supplementary Table S6.

**Figure 6.**
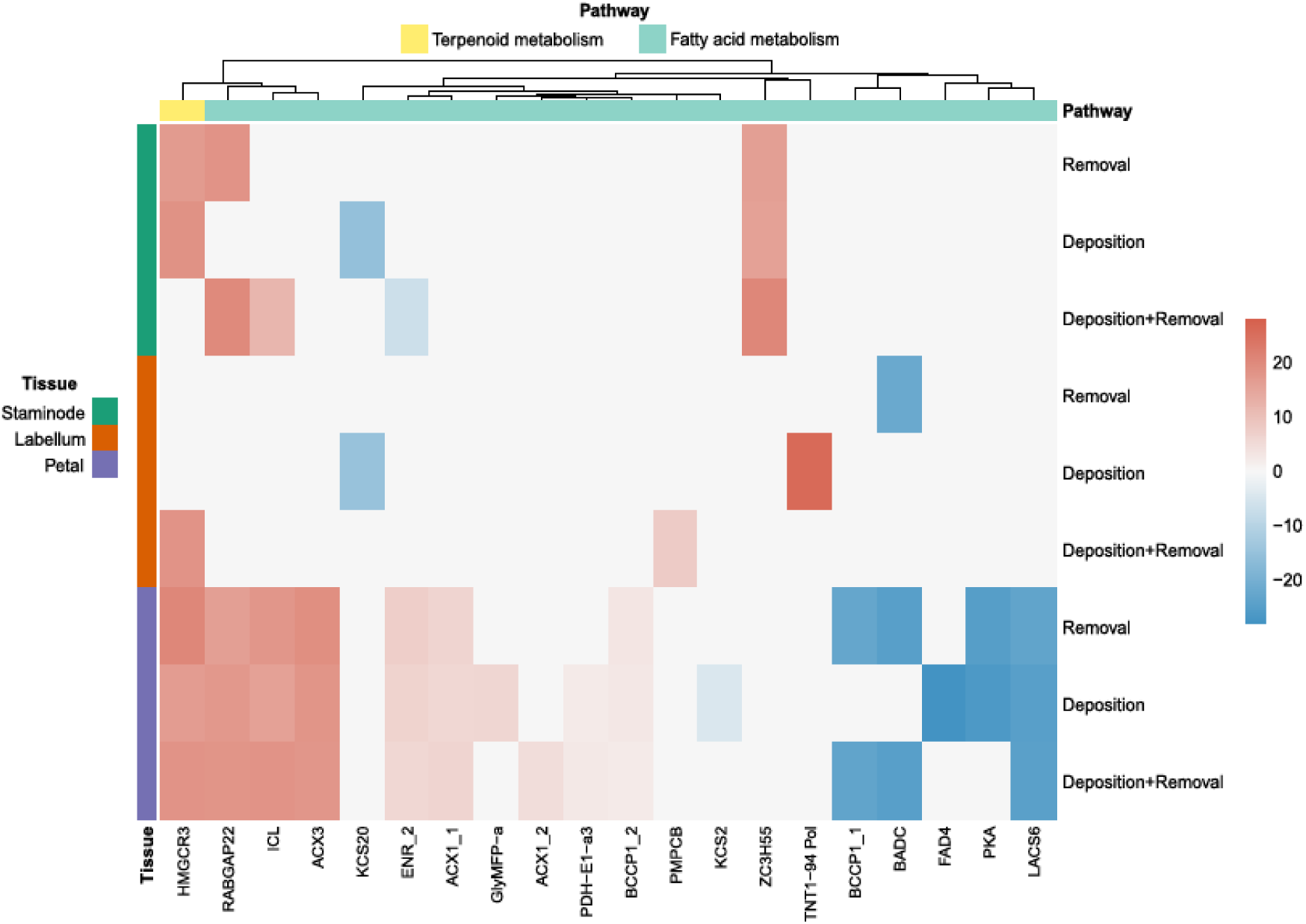
Heatmap showing differential expression of terpenoid and fatty acid metabolism genes of *C. henryi* in treatments compared to control. Colour intensities represent log_2_FC, with red indicating upregulation and blue indicating downregulation. *HMGCR3*: 3-hydroxy-3-methylglutaryl-coenzyme A reductase 3; *RABGAP22*: Rab GTPase-activating protein 22; *ICL*: Isocitrate Lyase; *ACX3*: Acyl-coenzyme A oxidase 3; *KCS20*: 3-ketoacyl-CoA synthase 20; *ENR*: Enoyl-[acyl-carrier-protein] reductase; *ACX1*: Acyl-coenzyme A oxidase 1; *GlyMFP-*α: Glyoxysomal fatty acid beta-oxidation multifunctional protein MFP-α; *PDH-E1-*α*3*: Pyruvate dehydrogenase E1 component subunit alpha-3; *BCCP1*: Biotin carboxylase 1; *PMPCB*: Mitochondrial-processing peptidase subunit beta; *KCS2*: 3-ketoacyl-CoA synthase 2; *ZC3H55*: Zinc finger CCCH domain-containing protein 55; *TNT1-94 Pol*: Retrovirus-related Pol polyprotein from transposon TNT 1-94; *BADC*: Biotin lipoyl attachment domain-containing protein; *FAD4*: Fatty acid desaturase 4; *PKA*: Pyruvate kinase isozyme A; *LACS6*: Long chain Acyl-CoA synthase 6.

## Discussion

### Floral scent diversity in the genus *Cypripedium*

#### Overview of floral scent in the genus Cypripedium

In the comparative study we discovered a substantial diversity of floral VOCs in *Cypripedium*. Across 31 sampled taxa and covering 24 out of the 54 accepted species of the genus, we identified 258 floral VOCs, predominantly terpenoids, aliphatics, and aromatics. While some floral VOCs were detected across multiple taxa, many were exclusive to a single taxon. To our knowledge, this study provides one of the most extensive genus-wide samplings of floral volatiles conducted thus far. The only broader genus-level analyses to date are those by Füssel *et al*. (2007) on 34 *Salix* species and Cao *et al*. (2026) on 32 *Ficus* species, both belonging to substantially larger genera (∼475 and ∼892 species, respectively; World Checklist of Selected Plant Families, accessed June 2026). Notably, our *Cypripedium* dataset exhibited greater chemical diversity than similarly sized datasets of the highly fragrant, euglossine bee-pollinated orchid genera *Stanhopea* (Williams and Whitten 1999), *Gongora* (Hetherington-Rauth and Ramírez 2016) and *Catasetum* (Milet-Pinheiro and Gerlach 2017). However, it is worth noting that aforementioned studies captured a smaller percentage of genus-wide variation, as these genera contain more species than *Cypripedium*. In our study, several taxa such as *C. californicum* and *C. calceolus* exhibited very complex floral scent profiles. While the number of individual VOCs varied substantially between species, the high cumulative diversity observed within *Cypripedium* may be driven by the broad range of pollinators exploited across the genus. The monoterpene alcohol linalool appears to be a central component of the *Cypripedium* chemical repertoire, particularly within section *Cypripedium* where it frequently dominates the scent bouquet. Such widespread occurrence aligns with the broader role of linalool as one of the most ubiquitous floral volatiles in angiosperms (Knudsen *et al.,* 2006; Farré-Armengol *et al.,* 2020), likely serving a dual function in these orchids as both a primary pollinator attractant and a defence against herbivory (L. Zhang *et al*., 2023) and floral pathogens (Shimada *et al.,* 2014).

#### Floral scent variation in Cypripedium: Pollinator influence and chemical relationships

Although pollinator group significantly influenced scent composition (p = 0.049), it accounted for only 10% of the total variance. This relatively low explanatory power aligns with a strategy of generalized food deception hypothesized for the majority of our sampled taxa. Unlike Batesian mimicry or sexually deceptive pheromone systems, generalized food deception lacks the stringent evolutionary pressure for high chemical specificity. The floral scents of taxa within the bees and floricolous flies pollination group contained several well-known bee attractants, including the aromatic methyl salicylate and the common terpenes linalool and (*E*)-β-ocimene (Williams and Whitten 1983; Gerlach and Schill 1991; Farré-Armengol *et al.,* 2017), which are likely associated with the presence of nectar and pollen by these insects. Electrophysiological studies support the functional relevance of these compounds to bees; linalool, in particular, has been shown to elicit strong antennal responses in known pollinator species of *Cypripedium*, specifically within the key pollinator genera *Lasioglossum*, *Andrena*, and *Apis* (Luo *et al.,* 2013; Braunschmid *et al.,* 2017). As linalool is a dominant constituent in numerous bee-pollinated lineages (Gong *et al.,* 2015; Krug *et al.,* 2018; Cordeiro and Dötterl 2023), its prevalence in our sampled taxa likely reflects an exploitation of conserved sensory biases in these hymenopterans. Among the three species in our dataset with documented bee and fly pollination, *C. calceolus* and *C. reginae* are associated with hoverflies, whereas *C. flavum* is visited by blowflies (Table S2). Interestingly, (*E*)-β-farnesene, an aphid alarm pheromone known to attract hoverflies in search of an oviposition site (Jiang *et al.,* 2020; B. Wang *et al.,* 2022), was present in our overall dataset but absent from the two hoverfly pollinated species *C. calceolus* and *C. reginae*. The absence of (*E*)-β-farnesene or other aphid-related VOCs in the floral fragrance of *C. calceolus* combined with the fact that the species is predominantly pollinated by generalist polylectic bees, suggests that hoverflies likely visit the flowers in search of nutritive rewards rather than oviposition sites. In contrast, *C. reginae* may indeed employ brood site mimicry, given that its most abundant floral VOC methyl salicylate (31%) and ethyl salicylate (7%) are well-known herbivore-induced plant volatiles (HIPVs). Methyl salicylate in particular, is frequently produced by plants in reaction to aphid infestation, which consequently exerts a strong attraction on hoverflies in search of an aphid colony for oviposition (Primante and Dötterl 2010; Rodriguez-Saona *et al.,* 2011; Salamanca *et al.,* 2019). Although a multimodal strategy integrating both visual and olfactory cues is likely, hoverflies often exhibit a strong reliance on visual floral cues (Laubertie *et al.,* 2006; Mishra *et al.,* 2025). This is exemplified by the tropical slipper orchid *Paphiopedilum barbigerum*, which is also hoverfly pollinated but lacks floral volatiles. In this species, the yellow-coloured staminode was hypothesized to be the primary visual attractant, leveraging the known attraction of hoverflies to this colour (Shi *et al.,* 2009). A comparable visual cue exists in *C. reginae*, whose staminode also exhibits distinct yellow spots. The bimodal volatile profile of *C. flavum* suggests an adaptive dual-signalling strategy adapted to the olfactory preferences of its pollinators, bumblebees and blowflies. The floral VOC δ-1-pyrroline and the two C5-branched compounds, which are associated with decay or fermentation odours, could increase the attractiveness to the pollinating blowfly species, while the presence of the terpenes (*Z/E*)-β-Ocimene, α-Copaene, and β-Caryophyllene, likely serves as the primary olfactory attractant for bumblebees (Dötterl and Gershenzon 2023). The drosophilid-pollinated *C. micranthum* and *C. bardolphianum* emitted scent profiles that were markedly different from the rest of the genus, characterized predominantly by aliphatic alcohols and a distinct lack of shared major VOCs. Although the fragrances of these two species were perceived as similarly sweet and alcoholic, their chemical compositions differed substantially. The floral scent of *C. bardolphianum* was defined by the aliphatic esters ethyl butyrate and ethyl propanoate, along with the phenolic compound 2-methoxyphenol. While the former two esters likely mimic the chemical signature of fermenting or overripe fruit attractive to most drosophilid species, 2-methoxyphenol is known to function not only as a floral attractant for moths (Knudsen *et al*., 1993) but also as a volatile marker of microbial (Tahir *et al*., 2019) and insect-driven (Revadi *et al*., 2021) degradation of plant tissues. In contrast, the profile of *C. micranthum* was dominated by the fermentation-associated aliphatic compounds acetoin and 2,3-butanediol. These latter compounds were also found in high concentrations in the drosophilid-pollinated *Aristolochia baetica* (Rupp *et al.,* 2024). The additional detection of β-cyclocitral, a known fruit fly attractant (Piñero *et al.,* 2019), provides further evidence for a highly specialized drosophilid-attraction mechanism in *C. micranthum*.

#### Phylogenetic signal in Cypripedium’s floral scent

Our analyses reveal a complex evolutionary pattern of floral scent variation within *Cypripedium*. At the sectional level, we did not detect a significant effect of phylogenetic proximity on floral fragrance similarity. This lack of significance may be attributed to the fact that taxonomical sections represent relatively coarse categories and high intra-sectional variability likely obscured scent-based differences between sections. Furthermore, several sections were represented by small sample sizes, partly due to our dataset composition and partly because sections such as *Flabellinervia* (n = 2), *Arietinum* (n = 2), and *Californica* (n = 1) are inherently species-poor. At the interspecific level however, the Mantel test revealed a significant, although weak, correlation between phylogenetic distance and floral scent dissimilarity (r=0.161). While this indicates the presence of a phylogenetic signal, the explanatory power was remarkably low (r^2^=0.026), suggesting that phylogenetic relatedness accounts for only ∼2.6% of the observed variation. This interpretation is further supported by the observation that even closely related species in our dataset often shared only a small number of floral scent compounds. A weak phylogenetic signal is consistent with the view of floral scent as a highly labile and rapidly evolving trait. Floral scent is often subject to intense selection by pollinators and local ecological conditions, which limits its usefulness for phylogenetic reconstruction (Williams and Whitten 1999; Levin *et al.,* 2003). The predominant role of ecology over ancestry is supported by our finding that, while sectional taxonomy had no effect, pollinator groups significantly influenced scent composition in *Cypripedium*. While phylogenetic relationships influence floral scent variation in some Orchidaceae lineages (Steiner *et al.,* 2011; Joffard *et al.,* 2020), the generalized food deception strategy employed by most *Cypripedium* taxa could contribute to the weak signal observed here. Because these orchids often target a broad pollinator spectrum, stabilizing selection on specific VOC profiles may be relaxed. This allows for high chemical plasticity and rapid divergence between species. The degree of phylogenetic signal in floral scent can be a product of pollinator specificity. For instance, Prieto Benítez *et al*. (2016) reported a stronger phylogenetic signal in nocturnal scents compared to diurnal scents in *Sileneae*, attributing this pattern to the more taxonomically constrained nature of the nocturnal pollinator assemblage compared to the diurnal one. Nevertheless, one must consider that distinct pollinator groups carry distinct evolutionary and sensory limitations. These pollinator constraints also directly influence the variability of floral scent bouquets over evolutionary time. To conclude, the broad and often overlapping pollinator assemblages across *Cypripedium* likely account for this pattern, where a significant but minimal phylogenetic signal is present but where ancestral constraints are secondary to the selective pressures of pollinator attraction, resulting in a scent profile that is functionally driven rather than phylogenetically conserved.

#### Intra-specific variation of floral scent in Cypripedium

High intraspecific variation was uncovered within the three sampled infraspecific taxa of both *C. macranthos* and *C. parviflorum*. In the latter, the variation was so prominent that the two sampled forms of var. *pubescens* shared only a single common floral VOC. Such chemical heterogeneity aligns with the polyphyletic nature of the var. *pubescens* complex recently identified by Lagou *et al*. (2024). These results provide further evidence of the systematic confusion surrounding the var. *pubescens* complex, highlighting the critical need for a taxonomic revision of this group. Variation in floral VOCs was also pronounced among the sampled infraspecific taxa of *C. macranthos*, but to a bit smaller amount. In the phylogenetic analysis by Lagou *et al*. (2024), which utilized a significantly more comprehensive sampling of *C. macranthos* infraspecific taxa than previous studies (Fatihah *et al.,* 2011; J. Li *et al.,* 2011; Szlachetko *et al.,* 2021), both *C. franchetii* and *C. yunnanense* were recovered as nested within the *C. macranthos* clade. While both species were included in the present study, our floral scent data neither contradict nor explicitly support this placement; neither species exhibited a profile particularly similar to the sampled varieties of *C. macranthos*. Specifically, *C. franchetii* shared only linalool, while *C. yunnanense* possessed no volatile organic compounds (VOCs) in common with the *C. macranthos* varieties. Interestingly, within this broader ‘*macranthos*’ clade, i.e., including *C. franchetii* and *C. yunnanense*, floral scent exhibited high functional lability even at shallow taxonomic scales. For instance, the scent profiles of *C. macranthos* var. *rebunense* and *C. macranthos* var. *hotei-atsumorianum* were the most similar (Figure 4), despite not being closely related within the species complex (Lagou *et al.,* 2024). Comparison of the floral scent profiles generated in this study with published volatile data for *Cypripedium* species revealed both expected similarities and several unexpected discrepancies. For *C. calceolus*, *C. reginae*, and *C. guttatum*, our samples largely confirm previously reported scent compositions, with our dominant core compounds matching published profiles and differences confined primarily to lower abundance VOCs (Bergstrom *et al.,* 1992; Barkman *et al.,* 1997; Braunschmid *et al.,* 2017, 2020). In contrast, published data for *C. parviflorum* var. *pubescens* already show marked variation. Bergström *et al*. (1992) and Barkman *et al*. (1997) reported strikingly different floral fragrance compositions, with the sample of the latter study lacking the otherwise highly dominant compound 1,3,5-trimethoxybenzene as well as most terpenoid VOCs. Our sample closely resembled the profile reported by Barkman *et al*. (1997), characterized by 2-phenylethanol as the major constituent, followed by benzaldehyde and phenylacetaldehyde. Surprisingly, the floral scent profiles we obtained for several species stands in stark contrast to previously published floral VOCs data, though similar methods were used among the studies. For instance, our findings failed to confirm the dominant compounds previously reported for *C. kentuckiense* (Barkman *et al*., 1997), *C. tibeticum* (P. Li *et al*., 2006), *C. plectrochilum* (P. Li *et al*., 2008a), *C. flavum* (Zheng *et al*., 2011), *C. micranthum*, and *C. bardolphianum* (P. Li, 2006). Most notably, in the cases of *C. tibeticum*, *C. flavum*, and *C. plectrochilum*, there was no overlap between the floral VOCs identified in our samples and the previously published profiles. Altogether, these observations demonstrate that floral VOC profiles in *Cypripedium* can be highly variable within species, suggesting that population-specific adaptive forces alongside genetic drift and gene flow shape floral VOC emission across a species’ range. Concurrently, the marked discrepancies between our findings and published data underscore a key methodological limitation of single-individual sampling. Fully capturing the spectrum of intraspecific chemical diversity across populations will require future studies to implement broader sampling regimes incorporating multiple individuals per species.

### Case study in *C. henryi*: post pollination effects on floral scent production

#### *Floral scent of* Cypripedium henryi

The floral scent of *C. henryi* was found to be strongly dominated by aliphatic compounds, which generally accounted for more than 90% of the total floral bouquet (Table 1). Analysis of the 24 floral scent samples revealed 33 VOCs, a substantially greater number than the four aliphatic volatiles (dodecanal, undecane, nonanal, and decanal) identified in the earlier study by Peng Li *et al*. (2008). Nonanal and decanal were not consistently identified as floral volatiles in our study, as they did not reach the required threshold of being at least fivefold more abundant in floral samples than in the corresponding leaf controls. Consistent with Peng Li *et al*. (2008), dodecanal and undecane emerged as the major components of the floral scent in our samples. The fatty aldehyde dodecanal was by far the most abundant compound in the floral fragrance of *C. henryi*. The presence of this compound, along with the co-detected C_12_ metabolic derivatives 1-dodecanol and dodecanoic acid, suggests a potential dual function beyond pollinator attraction, as these molecules are known to possess significant antimicrobial and antifungal activities (Togashi *et al.,* 2007; Davis Hanna *et al.,* 2008; Hall *et al.,* 2011; Giovagnoni *et al.,* 2022). Given that dodecanal was the heavily dominant compound in both our analysed samples and published literature, its high emission levels are unlikely to be linked to artifacts or stress responses induced by our experimental setup. Ultimately, this dual functionality of floral VOCs likely represents an evolutionary adaptation that simultaneously facilitates pollinator attraction and floral defence. Undecane was the second most prevalent compound, and although it has shown to act as an attractant for various types of insects (Y. Wang *et al.,* 2017; X. Zhang *et al.,* 2025) and more specifically as an alert pheromone for ants (Hölldobler and Wilson 1990; Xu *et al.,* 2023), its ecological role in *Cypripedium* pollination remains uncertain. The detection of terpenoid volatiles in the floral fragrance of *C. henryi* represents a new finding compared to previous studies. Specifically, the compounds linalool, 4-oxoisophorone, and 2,4,4-trimethyl-3-carboxaldehyde-5-hydroxy-2,5-cyclohexadien-1-one were consistently present across all floral samples analysed. Notably, the chemical composition of *C. henryi*’s floral scent differed greatly from that of its sister taxon *C. segawai*, which lacked aliphatic compounds and was dominated by terpenoids.

#### Effects of pollination on floral fragrance composition in C. henryi

Pollination treatment (pollinium removal and/or deposition) revealed only subtle effects on the floral scent of *C. henryi*, with the inter-individual variation between the two sampled plants contributing substantially to the observed variance (Figure 5). While we observed a significant reduction in total scent emission following cross-pollination (i.e. deposition treatment), the more pronounced 3.6-fold decrease observed in flowers of the deposition+removal treatment did not reach statistical significance. This difference in significance could very well be attributable to high inter-replicate variability rather than antagonistic physiological effects caused by the pollinia removal. A reduction in total scent emission represents the most widely documented post-pollination phenomenon in flowering plants. This response has been described across various Orchidaceae, including *Platanthera bifolia* (Tollsten 1993) and *Ophrys sphegodes* (Schiestl *et al.,* 1997), as well as in diverse taxa such as *Cirsium* spp (Theis and Raguso 2005) and *Silene latifolia* (Dötterl *et al.,* 2005; Muhlemann *et al.,* 2006). In *S. latifolia*, this downregulation is particularly robust, with scent emission decreasing four-fold (Muhlemann *et al.,* 2006) or even ceasing entirely (Dötterl *et al.,* 2005). In contrast, our results for *C. henryi* remain ambiguous due to high inter-replicate variability and a general, although non-significant, decline in volatile emissions across all groups. The general decrease in the absolute abundance of all major VOCs across all post-treatment samples suggests a broad senescence-related decline in *C. henryi* that may mask more subtle, specific post-pollination responses. Furthermore, we found no evidence of VOC upregulation acting as a pollinator repellent, a mechanism previously identified in *O. sphegodes* (Schiestl and Ayasse 2001). One prevailing assumption suggests that post-pollination changes are only initiated once pollen tubes reach the ovary, a process taking less than 48 hours in *Antirrhinum* and *Petunia* (Negre *et al.,* 2003), but which may vary significantly across species. While our results, alongside unpublished data from *C. calceolus* (S. Dötterl *et al*., unpublished data), might suggest that the genus *Cypripedium* lacks strong post-pollination scent cues, we caution against over-generalization. Post-pollination responses can vary significantly between congeneric species as it was found for *Cirsium* (Theis and Raguso 2005). Finally, we acknowledge the limitations imposed by our small sample size. Future studies with increased replicates, would be necessary to confirm if pollinated flowers truly exhibit a more substantial drop in total scent than unpollinated controls. Furthermore, given the late onset of scent production observed in *C. henryi*, we recommend delaying the initial sampling period to 72 hours post-anthesis.

#### Effects of pollination on floral volatile gene expression levels in C. henryi

Our results indicate that while transcriptional responses to pollination in *C. henryi* are detectable, they remain largely ambiguous. Expression profiles for FA and terpenoid biosynthetic genes lacked clear treatment-specific signatures; no identified genes exhibited uniform reactions across all tissue types, and almost none showed consistent patterns across the two treatments involving pollinia deposition. The only exception to this last point was the up-regulation of *PDH-E1-*α in petals. Despite its role in providing Acetyl-CoA for FA biosynthesis, its expression trajectory did not align with the volatile data. As FA-derived volatiles decreased across all groups and showed no treatment effect, the increased expression of *PDH-E1-*α was likely counterbalanced by downstream regulatory mechanisms or metabolic activity of other floral tissue. *BCCP1_1* and *BADC* displayed distinct localized responses being downregulated primarily in petals and only in treatments with pollinia removal (removal and deposition+removal). This pattern potentially reflects metabolic consequences of mechanical disturbance triggered by pollinia removal, independent of pollination. The parallel expression of these two genes is particularly notable given their antagonistic roles within the ACCase complex, the primary gatekeeper of FA biosynthesis (Page *et al*., 1994; Balbuena *et al*., 2025). While *BCCP1* provides the essential biotin-carrying subunit to drive the reaction, *BADC* acts as a competitive inhibitor to suppress it (Salie *et al*., 2016). Their synchronized downregulation could suggest a coordinated recalibration of the ACCase complex in response to pollinia removal. Furthermore, the divergent regulation of a second *BCCP1* isoform, which exhibited an antagonistic expression pattern to the first, underscores the sophisticated and multi-layered control of the ACCase complex in these tissues. *LACS6* encodes the enzyme responsible for activating free FAs into fatty acyl-CoA (Zhao *et al*., 2021), an obligatory step for their entry into peroxisomal β-oxidation pathway in which long-chain precursors are sequentially shortened to produce the aldehydes and alkanes that constitute FA-derived volatiles (Schwab *et al*., 2008). In our study, *LACS6* exhibited consistent down-regulation in petals of flowers having undergone a treatment, pointing to a coordinated, petal-specific shift in lipid metabolic balance. While a downregulation of *LACS6* offers a plausible molecular mechanism for a reduction in FA-derived volatiles (such as dodecanal and undecane), this transcriptional signal observed in treated flowers was not mirrored in the scent profiles. There was no statistical evidence nor any discernible trend in the scent data suggesting that absolute levels of *LACS6*-derived compounds were lower in treated flowers compared to controls. This divergence suggests that the metabolic shift may be more subtle than the transcriptional data implies, and may point to post-transcriptional regulations or compensatory pathways that stabilize volatile emission despite localized gene suppression. The only terpenoid-related gene identified through differential expression analysis was *HMGCR3,* which encodes the rate-limiting enzyme of the MVA pathway. This gene exhibited the most complex regulatory profile with its upregulation across treatments being highly tissue-specific (Figure 6). This upregulation is noteworthy since the MVA pathway is primarily associated with the synthesis of sesquiterpenes (Rohmer 1999; Lichtenthaler 1999), a class of compounds entirely absent from the floral scent of *C. henryi*. Consequently, these changes in MVA-pathway gene expression are probably linked to other metabolic functions, including sterol production or defence-related compounds, rather than to scent emission itself.

## Conclusion

Our study uncovered a high chemical heterogeneity in the floral fragrances across the genus *Cypripedium*, revealing a broad spectrum of scent bouquets that varied greatly in both the number and identity of floral volatiles. The pronounced interspecific variation revealed here, alongside existing volatile data, suggests that floral scent in this genus is a highly labile trait capable of rapid evolutionary divergence. Within this diversity, we uncovered a significant but weak phylogenetic signal in the floral scent profiles of the sampled dataset. On a broader scale, pollinator groups significantly explained some of the variance in floral scent composition, whereas taxonomic sections did not. This finding aligns with the genus’s primary strategy of generalized food deception, where the high lability of floral scents enables these deceptive orchids to break away from ancestral patterns and optimize their chemical profiles to exploit the pre-existing biases of local pollinator communities. The investigation into *C. henryi* provided limited evidence for a directed post-pollination effect on floral fragrance, as sample variation was explained far more by individual plant clone than by the pollination treatments. Instead, a general treatment-independent decline in floral volatiles across all flowers suggested a reaction induced by floral aging. While total scent emission did decrease significantly after cross-pollination only, this result should be interpreted with caution given the low statistical power and the fact that the finding was not confirmed in the treatment involving pollinia removal alongside cross-pollination. Furthermore, differential expression analysis of candidate genes related to volatile biosynthesis aligned with the volatile data in lacking clear treatment-specific signatures.

## List of Supplementary Data

Table S1. Overview of sampled *Cypripedium* taxa, sample sizes, material origin, sectional classifications and used gas chromatography (GC) split ratios.

Table S2. Summary of documented *Cypripedium* pollinators.

Table S3. Relative amounts (%) of floral volatiles detected in 31 taxa of *Cypripedium*.

Table S4. Unidentified compounds in floral scent of *Cypripedium*.

Table S5. Global summary of total significantly differentially expressed transcripts (padj < 0.05, ∣log2FC∣>1) across all treatment comparisons in *C. henryi*.

Table S6. Differentially expressed transcripts (DETs) of candidate genes involved in terpene and fatty acid biosynthetic pathways in *C. henryi*.

## Author contributions

Conceptualization: JBC, GK

Methodology: JBC, SP, SD, CS, EYH, GK

Resources and Supervision: GK, SP, SD

Formal analysis: JBC, SD, EYH, CS, SK

Investigation and Interpretation: JBC, SP, SD, CS, EYH, SK, GK

Writing - Original Draft: JBC, EYH, GK

Writing - Review & Editing: JBC, SP, SD, CS, EYH, SK, GK

## AI assistance acknowledgement

AI driven language processing tools were used during the drafting of this manuscript to assist with linguistic editing and the development of code for figure generation.

## Availability of Data and Materials

RNA data generated within this study will be deposited in the European Nucleotide Archive (ENA) at EMBL-EBI. The accession numbers will be provided prior to final publication.

## General acknowledgements

We are grateful to Sebastian Urban and his team for their horticultural assistance and maintenance of the *Cypripedium* collection.

