## Supplementary Data Tables (Table S1-S6)) for "Floral scent diversity and evolution in deceptive *Cypripedium,* and post-pollination effects on floral scent and its biosynthesis in *C. henryi*"

**Supplementary Material**

**Table S1. Overview of sampled *Cypripedium* taxa, sample sizes, material origin, sectional classifications and used gas chromatography (GC) split ratios.** Multiple samples per taxon were collected from the same individual. Taxa with n = 2 represent paired short (5 min) and long (15 min) collection durations. Where multiple split ratios are listed, the first value corresponds to the 5 min collection.

| **Taxa** | **Samples (n)** | **Origin** | **Section** | **GC split ratio** |
| --- | --- | --- | --- | --- |
| *C. bardolphianum* W.W.Sm. & Farrer | 2 | Hengduan Mountains Biotechnology Ltd., Chengdu, China | *Sinopedilum* | 1&10 |
| *C. calceolus* L. | 1 | Undocumented^*^ | *Cypripedium* | 3 |
| *C. californicum* A.Gray | 1 | Gartenwerkstatt Schreiner, Schwarzach am Main, Germany | *Californica* | 1 |
| *C. cordigerum* D.Don | 1 | Gartenwerkstatt Schreiner, Schwarzach am Main, Germany | *Cypripedium* | 5 |
| *C. farreri* W.W.Sm. | 1 | Hengduan Mountains Biotechnology Ltd., Chengdu, China | *Cypripedium* | 10 |
| *C. fasciolatum* Franch. | 2 | Undocumented | *Cypripedium* | 1 |
| *C. flavum* P.F.Hunt & Summerh. | 2 | Hengduan Mountains Biotechnology Ltd., Chengdu, China | *Obtusipetala* | 1&3 |
| *C. formosanum* Hayata | 2 | Undocumented^*^ | *Flabellinervia* | 3&1 |
| *C. franchetii* Rolfe | 2 | Undocumented^*^ | *Cypripedium* | 1&3 |
| *C. guttatum Sw.* | 1 | Undocumented^*^ | *Bifolia* | 1 |
| *C. froschii* Perner | 1 | Undocumented^*^ | *Cypripedium* | 1 |
| *C. henryi* Rolfe | 2 | Undocumented^*^ | *Cypripedium* | 1 |
| *C. japonicum* Thunb. | 1 | Hengduan Mountains Biotechnology Ltd., Chengdu, China | *Flabellinervia* | 1 |
| *C. kentuckiense* C.F.Reed | 2 | Undocumented^*^ | *Cypripedium* | 3&5 |
| *C. macranthos* var. *macranthos* Sw. | 2 | Undocumented^*^ | *Cypripedium* | 1 |
| *C. macranthos* var. *rebunense* Kudô | 1 | Gartenwerkstatt Schreiner, Schwarzach am Main, Germany | *Cypripedium* | 1 |
| *C. macranthos* var. *hotei-atsumorianum* Sadovski | 2 | Undocumented^*^ | *Cypripedium* | 1 |
| *C. micranthum* Franch. | 2 | Undocumented^*^ | *Sinopedilum* | 1&25 |
| *C. parviflorum* var. *makasin* (Farw.) Sheviak | 1 | Undocumented^*^ | *Cypripedium* | 1 |
| *C. parviflorum* var. *pubescence* f. *pubescence* (Willd.) O.W.Knight | 1 | Undocumented^*^ | *Cypripedium* | 1 |
| *C. parviflorum* var. *pubescence* f. *planipetalum* (Fernald) P.J.Cribb | 1 | Undocumented^*^ | *Cypripedium* | 1 |
| *C. passerinum* Richardson | 1 | Undocumented^*^ | *Obtusipetala* | 1 |
| *C. plectrochilum* Franch. | 2 | Hengduan Mountains Biotechnology Ltd., Chengdu, China | *Arietinum* | 3&1 |
| *C. reginae* Walter | 1 | Undocumented^*^ | *Obtusipetala* | 1 |
| *C. segawai* Masamune | 1 | Gartenwerkstatt Schreiner, Schwarzach am Main, Germany | *Cypripedium* | 1 |
| *C. tibeticum* King ex Rolfe | 2 | Hengduan Mountains Biotechnology Ltd., Chengdu, China | *Cypripedium* | 1 |
| *C. yatabeanum* Makino | 2 | Undocumented^*^ | *Bifolia* | 1&3 |
| *C. yunnanense* Franch. | 1 | Gartenwerkstatt Schreiner, Schwarzach am Main, Germany | *Cypripedium* | 5 |
| *C. x alaskanum* P.M.Br. | 1 | Undocumented^*^ | *Bifolia* | 1 |
| *C. x columbianum* Sheviak | 1 | Gartenwerkstatt Schreiner, Schwarzach am Main, Germany | *Cypripedium* | 3 |
| *C. x ventricosum* Sw. | 1 | Undocumented^*^ | *Cypripedium* | 1 |

**"Undocumented" refers to plant material that was already in cultivation at the Oberhof or the Munich-Nymphenburg Botanical Garden, respectively, prior to the establishment of the electronic accessioning system.*

**Table S2. Summary of documented *Cypripedium* pollinators.** Incorporating data from Pemberton (2013) alongside additional records identified in the literature review; these are marked with *.

| **Section** | **Species** | **Pollinator groups** | **Specific pollinators, key findings, Country of observation** | **Source** |
| --- | --- | --- | --- | --- |
| ***Cypripedium*** | *C. calceolus* | Bees, Hoverflies, Sawflies | Bees: *Andrena* spp*.* and *Lasioglossum* spp*.* (Germany, Sweden & Italy), *Halictus* spp*.* (Germany & Sweden), *Nomada* sp. (Germany & Denmark), *Colletes* sp*.* (Denmark).  Hoverflies: *Eristalis rupium, Pipiza austriaca, Platycheirus albimanus* (Germany).  Sawflies: *Hoplocampa plagiata* (Germany). | Nilsson (1979); Kull (2008); Antonelli et al. (2009); Braunschmid et al. (2017)*****  Braunschmid et al. (2020)***** |
|  | *C. yunnanense* | Sweat bees | *Lasioglossum zonulum euronotum* (China). | Bänziger et al. (2008) |
|  | *C. tibeticum* | Bumblebees | Large queens of *Bombus lepidus, B. lucorum, B. hypnorum*) are the effective pollinators; workers are frequent but ineffective due to size mismatch (China). | P. Li et al. (2006) |
|  | *C. macranthos* var. *rebunense* | Bumblebees | Queens of *Bombus pseudobaicalensis* (Japan). | Sugiura et al. (2001, 2002) |
|  | *C. calcicola* (*C. smithii*) | Bumblebees | Queens of *Bombus* spp. (China). | P. Li et al. (2006); Li and Luo (2009) |
|  | *C. henryi* | Sweat bees | females of *Lasioglossum sauterum* (China). | P. Li et al. (2008b) |
|  | *C. montanum* | Bees | *Andrena* spp*.*, *Halictus* spp., *Lasioglossum* spp*.*, *Ceratina* sp*.*, *Osmia* sp*.* (USA). | Bernhardt & Meier (2014) |
|  | *C. parviflorum* | Bees | Small miscellaneous bees. var. *pubescens* pollinated by *Osmia proxima, O. albiventris, O. subaustralis, Ceratina calcarata* (USA). | Curtis (1954); Stoutamire (1967); Light & MacConaill (2006); Herring (2007) |
|  | *C. shanxiense* | Bees and autopollination | *Lasioglossum zonulum*; delayed self-pollination is primary pollination mode (China). | Wang et al. (2025)***** |
|  | *C. candidum* | Bees | *Halictus* spp*., Lasioglossum* spp*., Andrena* sp*., Augochlorella* sp*.* (Canada). | Catling & Knerer (1980) |
|  | *C. kentuckiense* | Bees | *Anthophora abrupta* and *A. bomboides* are primary pollinators (USA). | Edens Meier et al. (2021)***** |
| ***Acaulia*** | *C. acaule* | Bumblebees | Queens of *Bombus vagans* and *B. ashtoni* (USA). | Stoutamire (1967); Primack & Stacy (1998); Davis (1986) |
| ***Arietinum*** | *C. plectrochilum* | Bees | *Lasioglossum* spp. and *Ceratina* sp. *Lasioglossum* better fits floral architecture (China). | P. Li et al. (2008a) |
|  | *C. arietinum* | Sweat bees | *Lasioglossum* sp. (subgenus *Dialictus*) (USA). | Stoutamire (1967) |
| ***Flabellinervia*** | *C. japonicum* | Bumblebees | Workers of *Bombus remotus*, *B. imitator* and occasionaly also *B. picipes* (China)  Queens of *B. ardens* and *B. diversus diversus* (Japan). | Sun et al. (2009); Suetsugu & Fukushima (2014)***** |
| ***Bifolia*** | *C. guttatum* | Sweat bees | *Lasioglossum virideglaucum, L. clypeinitens* and *L. sauterum.*  *L. mystaphium* almost certain pollinator and *L. allodalum* a probable one (China). | Bänziger et al. (2005) |
|  | *C. yatabeanum* | Unknown/Inferred | Introgression with *C. guttatum* suggests at least one common pollinator (likely *Lasioglossum* sp.). | Argue (2012)***** |
| ***Enantiopedilum*** | *C. fasciculatum* | Parasitic wasps | Females of *Cinetus* spp. (USA). | Ferguson & Donham (1999) |
| ***Sinopedilum*** | *C. micranthum* | Fruit flies | *Drosophila* spp. (China). | P. Li et al. (2012) |
|  | *C. bardolphianum* | Fruit flies | *Drosophila* spp. (China). | Zheng et al. (2010) |
| ***Trigonopedia*** | *C. sichuanense* | Dung flies | *Scathophaga* spp. (China). | P. Li et al. (2012) |
|  | *C. fargesii* | Hoverflies | *Cheilosia* sp. (China). | Ren et al. (2011); Ren et al. (2012) |
|  | *C. lichiangense* | Hoverflies | *Ferdinandea cuprea* (China). | Zheng et al. (2022)***** |
|  | *C. lentiginosum* | Hoverflies | *Ferdinandea formosana* (China). | Zhongjian et al. (2008)***** |
| ***Obtusipetala*** | *C. flavum* | Bees and Blowflies | Bees: *Bombus hypnorum, B. remotus* and *Andrena* spp.  Blowflies: *Calliphora vomitoria.*  Blowflies compared to Bumblebees less efficient pollinators but more frequent (China). | Zheng et al. (2011); Bänziger et al. (2008) |
|  | *C. reginae* | Leaf-cutter bees, Hoverflies | *Megachile melanophaea, M. centuncularis* and *Syrphus torvus* (USA). | Vogt (1990); Edens-Meier et al. (2011) |
|  | *C. passerinum* |  | Autogamous. | Catling (1990) |
| ***Subtropica*** | *C. subtropicum* | Hoverflies | *Episyrphus balteatus*, *Eristalinus arvorum*, *Dasysyrphus brunettii*, *Eupeodes confrater*, *Korinchia nova*, *Xanthandrus talamaui* (China). | Jiang et al. (2020)***** |
| ***Irapeana*** | *C. irapeanum* | Sweat bees | *Lasioglossum nyctere* is a floral visitor and possible pollinator (Mexico). | Lozano-Rodriguez et al. (2018)***** |

**Table S3. Relative amounts (%) of floral volatiles detected in 31 taxa of *Cypripedium*.** Data are based on single flower scent samples for each taxon and are arranged based on sections. *Cypripedium:* light orange*, Californica:* blue, *Arietinum:* light green, *Flabellinervia:* grey, *Sinopedilum:* violet, *Bifolia:* yellow, *Obtusipetala:* light blue. Volatiles identified through authentic standard are marked with an “s”. The Kovats retention index is indicated under Ri. Species abbreviations are the following: pm: *C. parviflorum* var. *makasin*, pp: *C. parviflorum* var. *pubescence* f. *pubescence*, ppp: *C. parviflorum* var. *pubescence* f. *planipetalum*, col: *C. colombianum*, ken: *C. kentuckiense*, hen: *C. henryi*, seg: *C. segawai*, cor: *C. corniculatum*, far: *C. farreri*, fas: *C. fasciolatum*, cal: *C. calceolus*, ven: *C. ventricosum*, ma: *C. macranthos* var. *macranthos*, mre: *C. macranthos* var. *rebunense*, mh: *C. macranthos* var. *hotei-atsumorianum*, fra: *C. franchetii*, yun: *C. yunnanense*, fro: *C. froschii*, tib: *C. tibeticum*, cali: *C. californicum,* ple: *C. plectrochilum*, for: *C. formosanum*, jap: *C. japonicum*, mic: *C. micranthus*, bar: *C. bardolphianum*, gut: *C. guttatum*, ala: *C. alaskanum*, yat: *C. yatabeanum*, reg: *C. reginae*, fla: *C. flavum*, pas: *C. passerinum*.

| Compounds | **Ri** | ***pm*** | ***pp*** | ***ppp*** | ***col*** | ***ken*** | ***hen*** | ***seg*** | ***cor*** | ***far*** | ***fas*** | ***cal*** | ***ven*** | ***ma*** | ***mre*** | ***mh*** | ***fra*** | ***yun*** | ***fro*** | ***tib*** | ***cali*** | ***ple*** | ***for*** | ***jap*** | ***mic*** | ***bar*** | ***gut*** | ***ala*** | ***yat*** | ***reg*** | ***fla*** | ***pas*** |
| --- | --- | --- | --- | --- | --- | --- | --- | --- | --- | --- | --- | --- | --- | --- | --- | --- | --- | --- | --- | --- | --- | --- | --- | --- | --- | --- | --- | --- | --- | --- | --- | --- |
| **Aliphatic compounds** |  |  |  |  |  |  |  |  |  |  |  |  |  |  |  |  |  |  |  |  |  |  |  |  |  |  |  |  |  |  |  |  |
| Acetoin^s^ | 702 | - | - | - | - | - | - | - | - | - | - | - | - | - | - | - | - | - | - | - | - | - | - | - | 80 | - | - | - | - | - | - | - |
| Ethyl propanoate^s^ | 706 | - | - | - | - | - | - | - | - | - | - | - | - | - | - | - | - | - | - | - | - | - | - | - | - | 13 | - | - | - | - | - | - |
| Methyl butyrate^s^ | 716 | - | - | - | - | - | - | - | - | - | - | - | - | - | - | - | - | - | - | - | - | - | - | - | - | 2 | - | - | - | - | - | - |
| (2*R*,3*R*)/(2*S*,3*S*)-2,3-Butanediol^s^ | 769 | - | - | - | - | - | - | - | - | - | - | - | - | - | - | - | - | - | - | - | - | - | - | - | 12 | - | - | - | - | - | - | - |
| *meso*-2,3-Butanediol^s^ | 782 | - | - | - | - | - | - | - | - | - | - | - | - | - | - | - | - | - | - | - | - | - | - | - | 2 | - | - | - | - | - | - | - |
| Ethyl butyrate^s^ | 797 | - | - | - | - | - | - | - | - | - | - | - | - | - | - | - | - | - | - | - | - | - | - | - | <1 | 48 | - | - | - | - | - | - |
| Methyl 2-oxobutyrate | 828 | - | - | - | - | - | - | - | - | - | - | - | - | - | - | - | - | - | - | - | - | - | - | - | - | <1 | - | - | - | - | - | - |
| (*Z*)-3-Hexen-1-ol^s^ | 852 | - | - | - | 1 | - | - | - | <1 | - | 3 | - | 2 | - | - | - | - | 23 | - | 56 | - | - | 36 | - | - | - | - | - | 47 | - | - | - |
| 1-Hexanol^s^ | 863 | - | - | - | 3 | - | 10 | - | 4 | - | - | - | <1 | - | - | - | - | - | - | - | - | - | - | - | - | - | <1 | <1 | - | - | 15 | - |
| Acetoin acetate^s^ | 886 | - | - | - | - | - | - | - | - | - | - | - | - | - | - | - | - | - | - | - | - | - | - | - | 1 | - | - | - | - | - | - | - |
| Propyl butyrate^s^ | 894 | - | - | - | - | - | - | - | - | - | - | - | - | - | - | - | - | - | - | - | - | - | - | - | - | <1 | - | - | - | - | - | - |
| Ethyl valerate^s^ | 896 | - | - | - | - | - | - | - | - | - | - | - | - | - | - | - | - | - | - | - | - | - | - | - | - | <1 | - | - | - | - | - | - |
| Pentyl acetate^s^ | 910 | - | - | - | - | - | - | - | - | - | - | <1 | - | - | - | - | - | - | - | - | - | - | - | - | - | - | - | - | - | - | - | - |
| Methyl hexanoate^s^ | 921 | - | - | - | - | - | - | - | - | - | - | - | - | - | - | - | - | - | - | - | - | - | - | - | - | - | <1 | - | - | 2 | - | - |
| 2,3-Butanediol monoacetate I | 922 | - | - | - | - | - | - | - | - | - | - | - | - | - | - | - | - | - | - | - | - | - | - | - | 1 | - | - | - | - | - | - | - |
| 2,3-Butanediol monoacetate II | 929 | - | - | - | - | - | - | - | - | - | - | - | - | - | - | - | - | - | - | - | - | - | - | - | <1 | - | - | - | - | - | - | - |
| Ethyl 3-hydroxybutanoate | 930 | - | - | - | - | - | - | - | - | - | - | - | - | - | - | - | - | - | - | - | - | - | - | - | - | <1 | - | - | - | - | - | - |
| Hexanoic acid^s^ | 970 | - | - | - | - | - | - | - | - | - | - | - | - | - | - | - | - | - | - | - | - | - | - | - | - | - | - | - | - | - | - | - |
| 3-Octanone^s^ | 984 | - | - | - | - | - | - | - | - | - | - | - | - | - | - | - | - | - | - | - | - | - | - | 68 | - | - | - | - | - | - | - | - |
| Methyl 2-hydroxy-4-methylpentanoate | 986 | - | - | - | - | - | - | - | - | - | - | - | - | - | - | - | - | - | - | - | - | - | - | - | - | - | - | - | - | 1 | - | - |
| 3-Octanol^s^ | 992 | - | - | - | - | - | - | - | - | - | - | - | - | - | - | - | - | - | - | - | - | - | 37 | 9 | <1 | <1 | - | - | - | - | 1 | - |
| Methyl 2-hydroxy-3-methylpentanoate | 992 | - | - | - | - | - | - | - | - | - | - | - | - | - | - | - | - | - | - | - | - | - | - | - | - | - | - | - | - | 16 | - | - |
| Ethyl hexanoate^s^ | 995 | - | - | - | - | - | - | - | - | - | - | - | - | - | - | - | - | - | - | - | - | - | - | - | <1 | <1 | - | - | - | - | - | - |
| (*Z*)-3-Hexenyl acetate^s^ | 1003 | - | - | - | - | - | - | - | - | - | - | <1 | - | - | - | - | - | - | - | - | - | - | - | - | - | - | - | - | - | - | - | - |
| Hexyl acetate^s^ | 1009 | - | - | - | - | - | - | - | - | - | - | 2 | <1 | - | - | - | - | - | - | - | - | - | - | - | <1 | - | - | - | - | - | - | - |
| Methyl heptanoate | 1021 | - | - | - | - | - | - | - | - | - | - | - | - | - | - | - | - | - | - | - | - | - | - | - | - | - | <1 | - | - | - | - | - |
| 1-Octanol^s^ | 1067 | - | - | - | - | - | - | - | 33 | 82 | 8 | - | - | - | - | - | - | - | - | - | - | - | - | - | - | - | - | - | - | - | - | - |
| (*Z*)-3-Nonenal^s^ | 1093 | <1 | - | - | - | - | - | - | - | - | - | - | - | - | - | - | - | - | - | - | - | - | - | - | - | - | - | <1 | - | - | - | - |
| Heptyl acetate^s^ | 1108 | - | - | - | - | - | - | - | - | - | - | <1 | <1 | - | - | - | - | - | - | - | - | - | - | - | - | - | - | - | - | - | - | - |
| Methyl octanoate^s^ | 1121 | - | - | - | - | - | - | - | <1 | - | - | - | - | - | - | - | - | - | - | - | - | - | - | - | - | - | 58 | 2 | - | - | - | - |
| Octyl formate | 1126 | - | - | - | - | - | - | - | <1 | - | - | - | - | - | - | - | - | - | - | - | - | - | - | - | - | - | - | - | - | - | - | - |
| Octanoic acid^s^ | 1159 | - | - | - | - | - | - | - | 1 | - | - | - | - | - | - | - | - | - | - | - | - | - | - | - | - | - | - | - | - | - | - | - |
| Methyl (2*E*)-2-octenoate | 1167 | - | - | - | - | - | - | - | - | - | - | - | - | - | - | - | - | - | - | - | - | - | - | - | - | - | <1 | - | - | - | - | - |
| 1-Nonanol^s^ | 1168 | - | - | - | - | - | - | - | <1 | - | - | - | - | - | - | - | - | - | - | - | - | - | - | - | - | - | - | - | - | - | - | - |
| Ethyl octanoate^s^ | 1193 | - | - | - | - | - | - | - | - | - | - | - | - | - | - | - | - | - | - | - | - | - | - | - | <1 | - | - | - | - | - | - | - |
| Octyl acetate^s^ | 1207 | - | - | - | - | - | - | - | <1 | - | <1 | 45 | 5 | - | - | - | - | - | - | - | - | - | - | - | - | - | - | - | - | - | - | - |
| a Methyl nonenoate | 1218 | - | - | - | - | - | - | - | - | - | - | - | - | - | - | - | - | - | - | - | - | - | - | - | - | - | <1 | - | - | - | - | - |
| Methyl nonanoate | 1220 | - | - | - | - | - | - | - | - | - | - | - | - | - | - | - | - | - | - | - | <1 | - | - | - | - | - | <1 | - | - | - | - | - |
| 1-Decanol^s^ | 1269 | - | - | - | - | - | - | - | 7 | 5 | 1 | <1 | 2 | - | - | - | - | - | - | - | - | - | - | - | - | - | - | - | - | - | - | - |
| 6-Tridecene^s^ | 1284 | - | - | - | - | - | - | - | - | - | 1 | - | <1 | - | - | - | - | - | - | - | - | - | - | - | - | - | - | - | - | - | - | - |
| (*Z*)-3-Nonenyl acetate^s^ | 1292 | - | - | - | <1 | - | - | - | - | - | - | <1 | - | - | - | - | - | - | - | - | - | - | - | - | - | - | - | - | - | - | - | - |
| a Methyl decenoate | 1304 | - | - | - | - | - | - | - | - | - | - | - | - | - | - | - | - | - | - | - | - | - | - | - | - | - | <1 | - | - | - | - | - |
| Nonyl acetate^s^ | 1306 | - | - | - | - | - | - | - | - | - | - | <1 | - | - | - | - | - | - | - | - | - | - | - | - | - | - | - | - | - | - | - | - |
| a Methyl decenoate | 1307 | - | - | - | - | - | - | - | - | - | - | - | - | - | - | - | - | - | - | - | - | - | - | - | - | - | 1 | - | - | - | - | - |
| a Methyl decadienoate | 1309 | - | - | - | - | - | - | - | - | - | - | - | - | - | - | - | - | - | - | - | - | - | - | - | - | - | <1 | - | - | - | - | - |
| Methyl decanoate | 1321 | - | - | - | - | - | - | - | - | - | 3 | - | - | - | - | - | - | - | - | - | - | - | - | - | - | - | 2 | - | - | - | - | - |
| Decanoic acid^s^ | 1356 | - | - | - | - | - | - | - | <1 | - | - | - | - | - | - | - | - | - | - | - | - | - | - | - | - | - | - | - | - | - | - | - |
| Ethyl (4*E*)-4-decenoate | 1377 | - | - | - | - | - | - | - | - | - | - | - | - | - | - | - | - | - | - | - | - | - | - | - | - | - | - | - | - | - | - | - |
| Hexyl hexanoate^s^ | 1383 | - | - | - | - | - | - | - | - | - | <1 | - | - | - | - | - | - | - | - | - | - | - | - | - | - | - | - | - | - | - | - | - |
| Octyl butanoate | 1386 | - | - | - | - | - | - | - | - | - | <1 | - | - | - | - | - | - | - | - | - | - | - | - | - | - | - | - | - | - | - | - | - |
| Ethyl decanoate^s^ | 1392 | - | - | - | - | - | - | - | - | - | 1 | - | - | - | - | - | - | - | - | - | - | - | - | - | <1 | - | - | - | - | - | - | - |
| Decyl acetate^s^ | 1406 | - | - | - | - | 9 | - | - | - | - | - | 20 | 2 | - | - | - | - | - | - | - | - | - | - | - | - | - | - | - | - | - | - | - |
| Dodecanal^s^ | 1409 | - | - | - | - | - | 47 | - | - | - | - | - | - | - | - | - | - | - | - | - | - | - | - | - | - | - | - | - | - | - | - | - |
| Methyl undecanoate | 1412 | - | - | - | - | - | - | - | - | - | <1 | - | - | - | - | - | - | - | - | - | - | - | - | - | - | - | - | - | - | - | - | - |
| 1-Dodecanol^s^ | 1471 | - | - | - | - | - | - | - | <1 | - | - | - | - | - | - | - | - | - | - | - | - | - | - | - | - | - | - | - | - | - | - | - |
| Pentadecane^s^ | 1496 | - | - | - | - | - | - | - | - | - | - | - | - | - | - | - | - | - | - | - | - | - | - | - | - | - | - | - | - | - | - | 63 |
| Methyl dodecanoate | 1521 | - | - | - | - | - | - | - | - | - | 59 | - | - | - | - | - | - | - | - | - | - | - | - | - | - | - | - | - | - | - | - | - |
| Octyl hexanoate | 1581 | - | - | - | - | - | - | - | - | - | <1 | - | - | - | - | - | - | - | - | - | - | - | - | - | - | - | - | - | - | - | - | - |
| Ethyl dodecanoate^s^ | 1591 | - | - | - | - | - | - | - | - | - | 12 | - | - | - | - | - | - | - | - | - | - | - | - | - | - | - | - | - | - | - | - | - |
| Dodecyl acetate^s^ | 1604 | - | - | - | - | - | - | - | - | - | - | <1 | - | - | - | - | - | - | - | - | - | - | - | - | - | - | - | - | - | - | - | - |
| (*Z*)-8-Heptadecene^s^ | 1677 | - | - | - | - | - | - | - | - | - | - | - | - | - | - | - | - | - | - | - | 2 | - | - | - | - | - | - | - | - | - | - | - |
| Heptadecane^s^ | 1696 | - | - | - | - | - | - | - | - | - | - | - | - | - | - | - | - | - | - | - | <1 | - | - | - | - | - | - | - | - | - | - | - |
| Ethyl tetradecanoate | 1789 | - | - | - | - | - | - | - | - | - | - | - | - | - | - | - | - | - | - | - | - | - | - | - | - | - | - | - | - | - | - | - |
| Methyl hexadecanoate | 1922 | - | - | 1 | - | - | - | - | - | - | - | - | - | - | - | - | - | - | - | - | - | - | - | - | - | - | - | - | - | - | - | - |
| **TOTAL** |  | **<1** | **-** | **1** | **5** | **9** | **58** | **-** | **46** | **88** | **92** | **69** | **15** | **-** | **-** | **-** | **-** | **23** | **-** | **56** | **3** | **-** | **73** | **76** | **97** | **65** | **65** | **3** | **47** | **20** | **16** | **63** |
| **Terpenoids** |  |  |  |  |  |  |  |  |  |  |  |  |  |  |  |  |  |  |  |  |  |  |  |  |  |  |  |  |  |  |  |  |
| 2-Methyl-2,5-divinyltetrahydrofuran | 915 | - | - | - | - | - | - | - | - | - | - | - | - | - | - | - | - | - | <1 | - | - | - | - | - | - | - | - | - | - | - | - | - |
| β-Myrcene^s^ | 990 | - | - | - | - | - | - | - | - | - | - | <1 | - | - | - | - | - | - | - | - | 4 | - | - | - | - | - | - | - | - | - | - | - |
| (*Z*)-β-Ocimene^s^ | 1036 | - | - | - | - | - | - | - | - | - | - | - | - | - | - | -<1 | - | - | - | - | 2 | - | - | - | - | - | - | - | - | - | 7 | - |
| Lavender lactone^s^ | 1043 | - | - | - | - | - | - | 6 | - | - | - | - | - | - | 7 | 22 | - | - | 15 | - | - | - | - | - | - | - | - | - | - | - | - | - |
| (*E*)-β-Ocimene^s^ | 1047 | - | - | - | - | 67 | - | - | - | - | - | <1 | <1 | - | - | <1 | - | - | <1 | - | 2 | - | - | - | - | - | - | - | - | - | 4 | - |
| Bergamal | 1054 | - | - | - | - | - | - | - | - | - | 1 | - | - | - | - | - | - | - | - | - | - | - | - | - | - | - | - | - | - | - | - | - |
| (*Z*)-Arbusculone^s^ | 1054 | - | - | - | - | - | - | 3 | - | - | - | - | - | - | 20 | 5 | - | - | 10 | - | - | - | - | - | - | - | - | - | - | - | - | - |
| 2-(1-Hydroxyethyl)-5-methyl-5-vinyl-tetrahydrofuran | 1062 | - | - | - | - | - | - | - | - | - | - | - | - | - | - | <1 | - | - | <1 | - | - | - | - | - | - | - | - | - | - | - | - | - |
| (*E*)-Arbusculone | 1072 | - | - | - | - | - | - | 7 | - | - | - | - | - | - | 4 | 2 | - | - | 5 | - | - | - | - | - | - | - | - | - | - | - | - | - |
| 2(1-Hydroxyethyl)-5-methyl-5-vinyl-tetrahydrofuran | 1075 | - | - | - | - | - | - | - | - | - | - | - | - | - | - | - | - | - | <1 | - | - | - | - | - | - | - | - | - | - | - | - | - |
| (*Z*)-Linalool oxide furanoid^s^ | 1076 | <1 | - | - | - | - | - | - | - | - | - | <1 | <1 | - | - | - | - | - | - | - | - | - | - | - | - | - | - | - | - | - | - | - |
| (*E*)-Linalool oxide furanoid^s^ | 1091 | 2 | - | - | - | - | - | - | - | - | - | <1 | <1 | - | - | - | - | - | - | - | - | - | - | - | - | - | - | - | - | - | - | - |
| Terpinolene^s^ | 1093 | - | - | - | - | - | - | - | - | - | - | - | - | - | - | - | - | - | - | - | <1 | - | - | - | - | - | - | - | - | - | - | - |
| Linalool^s^ | 1099 | 46 | - | - | - | - | 31 | 32 | - | - | - | 18 | 81 | 80 | 4 | 16 | 69 | - | - | - | 2 | - | - | - | - | - | - | - | - | - | - | - |
| *allo*-Ocimene^s^ | 1129 | - | - | - | - | - | - | - | - | - | - | - | <1 | - | - | - | - | - | <1 | - | <1 | - | - | - | - | - | - | - | - | - | - | - |
| Ocimene derivatives | 1132 | - | - | - | - | 17 | - | - | - | - | - | - | - | - | - | - | - | - | <1 | - | <1 | - | - | - | - | - | - | - | - | - | <1 | - |
| Epoxy-oxoisophorone^s^ | 1135 | - | - | - | - | - | - | - | - | - | - | - | - | - | - | - | - | - | 1 | - | - | 89 | - | - | - | - | - | - | - | - | - | - |
| *neoallo*-Ocimene^s^ | 1143 | - | - | - | - | - | - | - | - | - | - | - | - | - | - | - | - | - | - | - | <1 | - | - | - | - | - | - | - | - | - | - | - |
| Lilac aldehyde A^s^ | 1146 | - | - | - | - | - | - | 5 | - | - | - | - | - | - | 7 | 3 | - | - | 6 | - | - | - | - | - | - | - | - | - | - | - | - | - |
| 4-Oxoisophorone^s^ | 1148 | - | - | - | - | - | 11 | - | - | - | - | 1 | - | - | - | - | - | - | - | - | - | 11 | - | - | - | - | - | - | - | - | - | - |
| Lilac aldehyde B+C^s^ | 1157 | - | - | - | - | - | - | 10 | - | - | - | - | - | - | 7 | 4 | - | - | 8 | - | - | - | - | - | - | - | - | - | - | - | - | - |
| Lilac aldehyde D^s^ | 1170 | - | - | - | - | - | - | 6 | - | - | - | - | - | - | 2 | 2 | - | - | 3 | - | - | - | - | - | - | - | - | - | - | - | - | - |
| (*Z*)+(*E*)-Linalool oxide pyranoid^s^ | 1177 | 1 | - | - | - | - | - | - | - | - | - | <1 | - | - | - | - | - | - | - | - | - | - | - | - | - | - | - | - | - | - | - | - |
| 2,6-Dimethyl-3,7-octadiene-2,6-diol | 1188 | - | - | - | - | - | - | - | - | - | - | <1 | - | - | - | - | - | - | - | - | - | - | - | - | - | - | - | - | - | - | - | - |
| Lilac alcohol A^s^ | 1206 | - | - | - | - | - | - | <1 | - | - | - | - | - | - | 6 | 4 | - | - | 3 | - | - | - | - | - | - | - | - | - | - | - | - | - |
| Lilac alcohol B + C^s^ | 1216 | - | - | - | - | - | - | 6 | - | - | - | - | - | - | 10 | 7 | - | - | 7 | - | - | - | - | - | - | - | - | - | - | - | - | - |
| Verbenone^s^ | 1224 | - | - | - | - | - | - | <1 | - | - | - | - | - | - | - | - | - | - | <1 | - | <1 | - | - | - | - | - | - | - | - | - | - | - |
| β-Citronellol^s^ | 1226 | - | - | - | - | - | - | - | - | - | - | - | - | - | - | - | - | - | - | - | <1 | - | - | - | - | - | 2 | 1 | - | - | - | 4 |
| Nerol^s^ | 1229 | - | - | - | - | - | - | - | - | - | - | - | - | - | - | - | - | - | - | - | - | - | - | - | - | - | <1 | <1 | - | - | - | - |
| β-Cyclocitral^s^ | 1230 | - | - | - | - | - | - | - | - | - | - | - | - | 8 | - | - | - | - | - | 17 | - | - | - | - | 2 | - | - | - | - | - | - | - |
| Lilac alcohol D^s^ | 1231 | - | - | - | - | - | - | 10 | - | - | - | - | - | - | 2 | <1 | - | - | <1 | - | - | - | - | - | - | - | - | - | - | - | - | - |
| Neral^s^ | 1244 | - | - | - | - | - | - | - | - | - | - | - | - | - | - | - | - | - | - | - | <1 | - | - | - | - | - | - | - | - | - | - | - |
| Geraniol^s^ | 1253 | - | - | - | - | - | - | - | - | - | - | - | - | - | - | - | - | - | - | - | 3 | - | - | - | - | - | - | - | - | - | - | - |
| Methyl citronellate^s^ | 1258 | - | - | - | - | - | - | - | - | - | - | - | - | - | - | - | - | - | - | - | <1 | - | - | - | - | - | 1 | <1 | - | - | - | - |
| Geranial^s^ | 1272 | - | - | - | - | - | - | - | - | - | - | - | - | - | - | - | - | - | - | - | <1 | - | - | - | - | - | - | - | - | - | - | - |
| 3,7-Dimethyl-octa-1,7-dien-3,6-diol | 1272 | - | - | - | - | - | - | - | - | - | - | <1 | - | - | - | - | - | - | - | - | - | - | - | - | - | - | <1 | - | - | - | - | - |
| (*E*)-Linalool oxide acetate pyranoid | 1289 | - | - | - | - | - | - | - | - | - | - | <1 | - | - | - | - | - | - | - | - | - | - | - | - | - | - | - | - | - | - | - | - |
| Methyl geranate^s^ | 1323 | - | - | - | - | - | - | - | - | - | - | - | - | - | - | - | - | - | - | - | <1 | - | - | - | - | - | 2 | <1 | - | - | - | - |
| Lilac alcohol formate A^s^ | 1345 | - | - | - | - | - | - | - | - | - | - | <1 | - | - | - | - | - | - | - | - | - | - | - | - | - | - | - | - | - | - | - | - |
| 8-Oxo-linalool | 1347 | - | - | - | - | 7 | - | 4 | - | - | - | - | - | - | 2 | 2 | - | - | - | - | - | - | - | - | - | - | - | - | - | - | - | - |
| Lilac alcohol formate B+C^s^ | 1345 | - | - | - | - | - | - | - | - | - | - | <1 | - | - | - | <1 | - | - | - | - | - | - | - | - | - | - | - | - | - | - | - | - |
| Citronellyl acetate^s^ | 1350 | - | - | - | - | - | - | - | - | - | - | - | - | - | - | - | - | - | - | - | <1 | - | - | - | - | - | <1 | - | - | - | - | - |
| Lilac alcohol formate D^s^ | 1356 | - | - | - | - | - | - | <1 | - | - | - | <1 | - | - | - | <1 | - | - | - | - | - | - | - | - | - | - | - | - | - | - | - | - |
| Neryl acetate^s^ | 1362 | - | - | - | - | - | - | - | - | - | - | - | - | - | - | - | - | - | - | - | <1 | - | - | - | - | - | - | - | - | - | - | - |
| Geranyl acetate^s^ | 1380 | - | - | - | - | - | - | - | - | - | - | - | - | - | - | - | - | - | - | - | 6 | - | - | - | - | - | - | - | - | - | - | - |
| α-Copaene^s^ | 1392 | - | - | - | - | - | - | - | - | - | - | - | - | - | - | - | - | - | - | - | - | - | - | - | - | - | - | - | - | - | 1 | - |
| β-Elemene | 1405 | - | - | - | - | - | - | - | <1 | - | - | - | - | - | - | - | - | - | - | - | - | - | - | - | - | - | - | - | - | - | - | - |
| Isocaryophyllene^s^ | 1426 | - | - | - | <1 | - | - | - | <1 | - | - | - | - | - | - | - | - | - | - | - | - | - | - | - | - | - | - | <1 | - | - | - | - |
| Neryl acetone^s^ | 1435 | - | - | - | - | - | - | - | <1 | - | - | - | - | - | - | - | - | - | - | - | 1 | - | - | - | - | - | - | - | - | - | - | - |
| β-Caryophyllene^s^ | 1442 | - | - | - | 23 | - | - | - | 2 | - | - | - | - | - | - | - | - | - | - | - | - | - | - | - | - | - | <1 | 1 | - | - | 1 | - |
| Dihydro-ß-ionone^s^ | 1450 | - | - | - | - | - | - | - | - | - | - | - | - | 3 | - | - | - | - | - | - | - | - | - | - | <1 | - | - | <1 | 53 | - | - | - |
| Geranyl acetone^s^ | 1454 | - | - | - | - | - | - | - | 6 | - | - | - | - | - | - | - | - | - | - | - | 15 | - | - | - | - | - | - | - | - | - | - | - |
| (*E*)-β-Farnesene^s^ | 1458 | - | - | - | - | - | - | - | <1 | - | <1 | - | - | - | - | - | - | - | - | - | <1 | - | - | - | - | - | - | - | - | - | - | - |
| α-Caryophyllene^s^ | 1476 | - | - | - | 3 | - | - | - | <1 | - | - | - | - | - | - | - | - | - | - | - | - | - | - | - | - | - | <1 | <1 | - | - | <1 | - |
| (*Z*,*Z*)-α-Farnesene^s^ | 1495 | - | - | - | - | - | - | - | <1 | - | 1 | - | - | - | - | - | - | - | - | - | - | - | - | - | - | - | - | - | - | - | - | - |
| β-Ionone^s^ | 1497 | - | - | - | - | - | - | - | - | - | - | - | - | 9 | - | - | - | - | - | - | - | - | - | - | - | - | - | - | - | - | - | - |
| Germacrene D^s^ | 1502 | - | - | - | - | - | - | - | - | - | - | - | - | - | - | - | - | - | - | - | - | - | - | - | - | - | - | - | - | - | <1 | - |
| (*E*,*E*)-α-Farnesene^s^ | 1510 | - | - | - | - | - | - | - | <1 | - | - | <1 | - | - | - | - | - | - | - | - | <1 | - | - | - | - | - | - | - | - | - | - | - |
| (*E*)-Nerolidol^s^ | 1568 | <1 | - | - | - | - | - | - | - | - | - | - | - | - | <1 | <1 | - | - | - | - | - | - | - | - | - | - | - | - | - | - | - | - |
| (*E*,*E*)-Farnesol^s^ | 1726 | - | - | - | - | - | - | - | <1 | - | - | - | - | - | - | - | - | - | - | - | 2 | - | - | - | - | - | - | - | - | - | - | - |
| (*E*,*E*)-Farnesal^s^ | 1750 | - | - | - | - | - | - | - | <1 | - | - | - | - | - | - | - | - | - | - | - | <1 | - | - | - | - | - | - | - | - | - | - | - |
| (*E*,*E*)-Methyl farnesoate | 1788 | - | - | - | - | - | - | - | - | - | - | - | - | - | - | - | - | - | - | - | <1 | - | - | - | - | - | - | - | - | - | - | - |
| Hexahydrofarnesylacetone^s^ | 1845 | - | - | 2 | - | - | - | - | - | - | - | - | - | - | - | - | - | - | - | - | - | - | - | - | - | - | - | - | - | - | - | - |
| **TOTAL** |  | **51** | **-** | **2** | **26** | **91** | **42** | **90** | **8** | **-** | **5** | **22** | **83** | **100** | **73** | **70** | **69** | **-** | **65** | **17** | **41** | **100** | **-** | **-** | **2** | **-** | **7** | **4** | **53** | **-** | **14** | **4** |
| **Aromatic compounds** |  |  |  |  |  |  |  |  |  |  |  |  |  |  |  |  |  |  |  |  |  |  |  |  |  |  |  |  |  |  |  |  |
| Benzaldehyde^s^ | 963 | - | 18 | 26 | - | - | - | - | 2 | - | - | - | - | - | - | - | - | - | - | - | 14 | - | - | - | - | - | - | 15 | - | - | - | - |
| Methoxymethylbenzene | 990 | - | - | - | - | - | - | - | - | - | - | - | - | - | - | - | - | - | - | - | 4 | - | - | - | - | - | - | - | - | - | - | - |
| 4-Methylanisole^s^ | 1023 | - | - | - | - | - | - | - | - | - | - | - | - | - | - | - | - | - | - | - | 2 | - | 27 | - | - | - | - | - | - | - | - | - |
| Benzyl alcohol^s^ | 1036 | - | 1 | - | - | - | - | - | 44 | - | - | 1 | - | - | - | - | - | - | - | - | 25 | - | - | - | - | 2 | 21 | 78 | - | - | - | - |
| Phenylacetaldehyde^s^ | 1047 | - | 3 | - | 1 | - | - | - | - | - | - | - | - | - | - | - | - | - | - | - | - | - | - | - | - | - | - | - | - | - | - | - |
| Salicylaldehyde | 1048 | - | - | - | - | - | - | - | - | - | - | - | - | - | - | - | - | - | - | - | - | - | - | - | - | - | - | - | - | - | 6 | - |
| Benzyl formate^s^ | 1079 | - | - | - | - | - | - | - | <1 | - | - | <1 | - | - | - | - | - | - | - | - | - | - | - | - | - | <1 | - | <1 | - | - | - | - |
| 2-Methoxyphenol^s^ | 1092 | - | - | - | - | - | - | - | - | - | - | - | - | - | - | - | - | - | - | - | - | - | - | - | - | 24 | - | - | - | - | - | - |
| Methyl benzoate^s^ | 1098 | - | - | - | - | - | - | - | - | - | - | - | - | - | - | - | - | - | - | - | - | - | - | - | - | - | - | - | - | 19 | - | - |
| 2-Phenylethanol^s^ | 1118 | - | 74 | - | 31 | - | - | - | - | 12 | - | <1 | - | - | - | - | - | 38 | - | - | <1 | - | - | - | - | - | - | - | - | - | - | - |
| Benzyl acetate^s^ | 1166 | - | - | - | - | - | - | - | <1 | - | - | 2 | - | - | - | - | - | - | - | - | - | - | - | - | - | - | - | <1 | - | - | - | - |
| 1,4-Dimethoxybenzene^s^ | 1166 | 32 | - | 59 | - | - | - | - | - | - | - | - | - | - | - | - | - | - | - | - | 11 | - | - | - | - | - | 4 | - | - | - | - | - |
| Ethyl benzoate | 1174 | - | - | - | - | - | - | - | - | - | - | - | - | - | - | - | - | - | - | - | - | - | - | - | - | <1 | - | - | - | 17 | - | - |
| 2-Phenethyl formate^s^ | 1181 | - | 2 | - | - | - | - | - | - | - | - | - | - | - | - | - | - | - | - | - | - | - | - | - | - | - | - | - | - | - | - | - |
| Methyl salicylate^s^ | 1202 | - | - | - | - | - | - | - | - | - | - | - | - | - | - | - | - | - | - | - | - | - | - | - | - | - | - | - | - | 31 | - | - |
| 1-Phenylbutane-2,3-dione^s^ | 1211 | - | - | - | - | - | - | - | - | - | - | - | - | - | - | - | - | 34 | - | - | - | - | - | - | - | - | - | - | - | - | - | - |
| 3-Phenyl-1-propanol^s^ | 1234 | - | - | - | - | - | - | - | - | - | - | - | - | - | - | - | - | - | - | - | - | - | - | - | - | <1 | - | - | - | - | - | - |
| 2-Phenyletyl acetate^s^ | 1260 | - | - | - | 34 | - | - | - | - | - | - | 5 | - | - | - | - | - | - | - | - | - | - | - | - | <1 | - | - | - | - | - | - | - |
| *p*-Anisaldehyde^s^ | 1263 | 12 | - | - | - | - | - | - | - | - | - | - | - | - | - | - | - | - | - | - | - | - | - | 17 | - | - | - | - | - | - | - | - |
| Ethyl salicylate | 1278 | - | - | - | - | - | - | - | - | - | - | - | - | - | - | - | - | - | - | - | - | - | - | - | - | <1 | - | - | - | 7 | - | - |
| (*E*)-Cinnamaldehyde^s^ | 1278 | - | - | - | - | - | - | - | - | - | - | - | - | - | - | - | - | - | - | - | - | - | - | - | - | 1 | - | - | - | - | - | - |
| *p*-Anisyl alcohol | 1287 | - | - | - | - | - | - | - | - | - | - | - | - | - | - | - | - | - | - | - | - | - | - | 7 | - | - | - | - | - | - | - | - |
| (*E*)-Cinnamyl alcohol^s^ | 1311 | - | - | - | - | - | - | - | - | - | - | - | - | - | - | - | - | - | - | - | - | - | - | - | - | <1 | - | - | - | - | - | - |
| Benzyl butyrate^s^ | 1350 | - | - | <1 | - | - | - | - | - | - | - | - | - | - | - | - | - | - | - | - | - | - | - | - | - | - | - | - | - | - | - | - |
| Eugenol^s^ | 1364 | - | - | - | - | - | - | - | - | - | - | <1 | - | - | - | - | - | - | - | - | - | - | - | - | - | - | - | - | - | - | - | - |
| 1,2,4-Trimethoxybezene^s^ | 1372 | - | - | - | - | - | - | - | - | - | - | - | - | - | - | - | 31 | - | - | 8 | - | - | - | - | - | - | - | - | - | - | - | - |
| Butyl benzoate | 1379 | - | - | - | - | - | - | - | - | - | - | - | - | - | - | - | - | - | - | - | - | - | - | - | - | <1 | - | - | - | - | - | - |
| Methyleugenol^s^ | 1404 | - | - | - | - | - | - | - | - | - | - | - | - | - | - | - | - | - | - | - | <1 | - | - | - | - | - | 3 | <1 | - | - | - | 4 |
| 2- or 3-Methylbutyl benzoate | 1446 | - | - | - | - | - | - | - | - | - | - | - | - | - | - | - | - | - | - | - | - | - | - | - | - | <1 | - | - | - | - | - | - |
| 2-Phenylethyl butanoate | 1448 | - | - | - | <1 | - | - | - | - | - | - | - | - | - | - | - | - | - | - | - | - | - | - | - | - | - | - | - | - | - | - | - |
| Pentyl benzoate | 1482 | - | - | - | - | - | - | - | - | - | - | - | - | - | - | - | - | - | - | - | - | - | - | - | - | - | - | - | - | <1 | - | - |
| cf 1,4-Dimethylindanyl actetate | 1666 | - | - | 4 | <1 | - | - | <1 | - | - | - | <1 | <1 | - | - | - | - | - | - | - | - | - | - | - | - | - | - | - | - | - | - | - |
| Methyl eudesmate | 1727 | - | - | - | <1 | - | - | - | - | - | - | - | - | - | - | - | - | - | - | - | - | - | - | - | - | - | - | - | - | - | - | - |
| Benzyl benzoate^s^ | 1784 | - | - | 7 | - | - | - | - | - | - | - | - | - | - | - | - | - | - | - | - | - | - | - | - | - | - | - | - | - | 1 | - | - |
| **TOTAL** |  | **44** | **98** | **96** | **68** | **-** | **-** | **<1** | **45** | **12** | **-** | **9** | **<1** | **-** | **-** | **-** | **31** | **72** | **-** | **8** | **56** | **-** | **27** | **24** | **<1** | **30** | **28** | **93** | **-** | **75** | **6** | **4** |
| **C5-Branched Chain compounds** |  |  |  |  |  |  |  |  |  |  |  |  |  |  |  |  |  |  |  |  |  |  |  |  |  |  |  |  |  |  |  |  |
| 3-methyl-1-butanol^s^ | 728 | - | - | - | - | - | - | - | - | - | - | - | - | - | - | - | - | - | - | - | - | - | - | - | <1 | - | - | - | - | - | - | - |
| 2-methyl-1-butanol^s^ | 730 | - | - | - | - | - | - | - | - | - | - | - | - | - | - | - | - | - | - | - | - | - | - | - | <1 | - | - | - | - | - | 7 | - |
| 3-Methyl-2-butenal | 779 | - | - | - | - | - | - | - | - | - | - | - | - | - | - | - | - | - | - | - | - | - | - | - | - | - | - | - | - | - | 4 | - |
| Ethyl 2-methylbutyrate | 846 | - | - | - | - | - | - | - | - | - | - | - | - | - | - | - | - | - | - | - | - | - | - | - | - | <1 | - | - | - | - | - | - |
| Methyl tiglate^s^ | 863 | - | - | - | - | - | - | - | - | - | - | - | - | - | - | - | - | - | - | - | - | - | - | - | - | <1 | - | - | - | - | - | - |
| 3-Methylbutyl acetate^s^ | 872 | - | - | - | - | - | - | - | - | - | - | - | - | - | - | - | - | - | - | - | - | - | - | - | <1 | - | - | - | - | - | - | - |
| Methyl 2-hydroxy-3-methylbutanoate | 890 | - | - | - | - | - | - | - | - | - | - | - | - | - | - | - | - | - | - | - | - | - | - | - | - | - | - | - | - | 4 | - | - |
| Ethyl tiglate | 936 | - | - | - | - | - | - | - | - | - | - | - | - | - | - | - | - | - | - | - | - | - | - | - | <1 | 4 | - | - | - | - | - | - |
| **TOTAL** |  | **-** | **-** | **-** | **-** | **-** | **-** | **-** | **-** | **-** | **-** | **-** | **-** | **-** | **-** | **-** | **-** | **-** | **-** | **-** | **-** | **-** | **-** | **-** | **<1** | **5** | **-** | **-** | **-** | **4** | **11** | **-** |
| **N-bearing compounds** |  |  |  |  |  |  |  |  |  |  |  |  |  |  |  |  |  |  |  |  |  |  |  |  |  |  |  |  |  |  |  |  |
| δ-1-Pyrroline | 674 | - | - | - | - | - | - | - | - | - | - | - | - | - | - | - | - | - | - | - | - | - | - | - | - | - | - | - | - | - | 15 | - |
| Phenylacetonitrile^s^ | 1143 | - | - | - | - | - | - | - | - | - | - | - | <1 | - | 3 | - | - | - | - | - | - | - | - | - | - | - | - | - | - | - | - | - |
| Indole^s^ | 1303 | 5 | 2 | - | 1 | - | - | - | - | - | - | - | - | - | - | - | - | - | - | - | - | - | - | - | - | - | - | - | - | - | - | - |
| **TOTAL** |  | **5** | **2** | **-** | **1** | **-** | **-** | **-** | **-** | **-** | **-** | **-** | **<1** | **-** | **3** | **-** | **-** | **-** | **-** | **-** | **-** | **-** | **-** | **-** | **-** | **-** | **-** | **-** | **-** | **-** | **15** | **-** |
| **Miscellaneous cyclic compounds** |  |  |  |  |  |  |  |  |  |  |  |  |  |  |  |  |  |  |  |  |  |  |  |  |  |  |  |  |  |  |  |  |
| Folifolone | 1109 | - | - | - | - | - | - | - | - | - | - | - | - | - | - | - | - | - | - | - | <1 | - | - | - | - | - | - | - | - | - | - | - |
| **TOTAL** |  | **-** | **-** | **-** | **-** | **-** | **-** | **-** | **-** | **-** | **-** | **-** | **-** | **-** | **-** | **-** | **-** | **-** | **-** | **-** | **<1** | **-** | **-** | **-** | **-** | **-** | **-** | **-** | **-** | **-** | **-** | **-** |
| **Spiroacetals** |  |  |  |  |  |  |  |  |  |  |  |  |  |  |  |  |  |  |  |  |  |  |  |  |  |  |  |  |  |  |  |  |
| 2-Methyl-1,7-dioxaspiro (5,5)-undecane^s^ | 1145 | - | - | - | - | - | - | - | - | - | - | - | - | - | - | - | - | - | - | - | - | - | - | - | <1 | - | - | - | - | - | - | - |
| **TOTAL** |  | **-** | **-** | **-** | **-** | **-** | **-** | **-** | **-** | **-** | **-** | **-** | **-** | **-** | **-** | **-** | **-** | **-** | **-** | **-** | **-** | **-** | **-** | **-** | **<1** | **-** | **-** | **-** | **-** | **-** | **-** | **-** |
| **TOTAL identified** |  | **100** | **100** | **100** | **100** | **100** | **100** | **90** | **100** | **100** | **97** | **99** | **100** | **100** | **75** | **70** | **100** | **94** | **65** | **81** | **99** | **100** | **100** | **100** | **100** | **99** | **99** | **100** | **100** | **100** | **63** | **71** |

**Table S4. Unidentified compounds in floral scent of *Cypripedium*.** ID of the compounds contain their respective Kovats retention index. Same species abbreviation used as in Table S3.

| **ID** | **Mass spectrum** | **Species** |
| --- | --- | --- |
| unk_739 | (39,41,53,67,79,81,95,110) | *cal, fro, mre, mh* |
| unk_746 | (43,53,71,80) | *fla* |
| unk_839 | (39,41,53,67,95,96) | *seg, fro, mh* |
| unk_848 | (40,43,44,45,54,71,77,98,121,131) | *mic* |
| unk_850 | (43,55,77,79,121,151) | *mic* |
| unk_942 | (40,45,57,69,74,85) | *mic* |
| unk_947 | (40,43,57,69,87,97,193) | *mic* |
| unk_967 | (40,56,69,84,99,112) | *fla* |
| unk_997 | (40,43,57,58,71,85,105,114,120) | *pas* |
| unk_1015 | (41,55,69,79,81,91,107,122,135,150) | *cal* |
| unk_1026 | (41,57,68,83,97) | *fla* |
| unk_1053 | (69,41,82) | *cal* |
| unk_1059 | (40,53,55,57,71,79,81,109,138) | *fro* |
| unk_1063 | (41,56,85) | *fla* |
| unk_1067 | (39,43,53,67,79,81,110) | *seg, fro, mh* |
| unk_1075 | (43,55,69,83,93,111,125,141) | *mh* |
| unk_1079a | (40,43,55,71,85,97,124,135) | *mh* |
| unk_1079b | (40,51,65,79,91,107,121,135,150) | *cali* |
| unk_1093 | (41,55,69,91,93,105,117,121,132) | *ven* |
| unk_1097 | (43,53,79,81,109,152) | *fro, mre, mh* |
| unk_1113 | (40,55,67,68,81,96,109,124) | *seg, fro, mh* |
| unk_1114 | (40,57,71,85,97,126) | *fla* |
| unk_1119 | (40,41,56,67,99,120,135) | *fla* |
| unk_1122 | (101,83) | *bar* |
| unk_1129 | (40,57,103,118) | *bar* |
| unk_1136 | (40,66,79,94,122,138) | *seg, mh* |
| unk_1149 | (40/41,55,67,81,96,109,121,123,138) | *mh* |
| unk_1162 | (43,57,71,85,96,126,127) | *mre, tib, mh* |
| unk_1210 | (43,55,67,77,93,111,121,136) | *seg, mh* |
| unk_1225 | (80,149) | *fla* |
| unk_1232 | (58,100,86) | *fla* |
| unk_1240 | (43,55,71,81,93,111,150,169) | *mre, seg* |
| unk_1261 | (43,55,71,81,93,111,123,138,169) | *seg* |
| unk_1263 | (43,55,81,93,109,123,138) | *seg, mre, mh* |
| unk_1269a | (43, 55,69,81,93,109,123,138) | *fro, mre* |
| unk_1269b | (43,55,81,93,109,123,138) | *seg* |
| unk_1271 | (43,57,71,85,112,113,327) | *mre* |
| unk_1273 | (40,55,69,81,93,111,121,137,152,168,183) | *fro* |
| unk_1283 | (40,43,55,71,96,111,135,150) | *fro, seg* |
| unk_1287 | (40,55,67,93,111,136,183) | *seg* |
| unk_1292 | (39,51,65,91,120) | *fro, mre, seg, mh* |
| unk_1297 | (40,43,57,71,91,119) | *seg* |
| unk_1302 | (43,57,67,81,82,96,109,123,138) | *fro, seg, mh* |
| unk_1335 | (43,55,71,82,98,121,135) | *fro, seg* |
| unk_1351 | (39,51,77,91,106,134) | *mh* |
| unk_1360 | (111,93) | *cal* |
| unk_1362 | (41,55,77,81,91,105,119,161,204) | *fla* |
| unk_1369 | (40,55,69,87,110,113,141,152,176) | *gut* |
| unk_1382 | (43,55,67,95,123,137) | *mre, seg, mh* |
| unk_1392 | (40,55,67,85,98,119,151,166) | *mre, seg, mh* |
| unk_1393 | (40,55,73,81,111,113,154,182,327) | *gut* |
| unk_1405 | (41,55,79,81,93,105,119,133,147,161,189,204) | *fla* |
| unk_1410 | (43,55,79,94,107,122) | *mre, seg, mh* |
| unk_1415 | (43,55,68,79,94,122) | *seg* |
| unk_1425 | (41,57,79,91,107,121,135,150) | *cali* |
| unk_1433 | (41,43,55,74,83,87,111,115,143,155) | *fas, gut, pas* |
| unk_1438 | (41,55,79,91,105,120,133,147,161,204) | *fla* |
| unk_1445 | (43,65,91,119,162) | *yun* |
| unk_1448 | (43,55,67,82,110,119,137,152) | *cal* |
| unk_1449 | (41,55,79,91,105,119,133,147,161,204) | *fla* |
| unk_1465 | (41,55,81,91,105,119,133,161,204) | *fla* |
| unk_1484 | (40,55,69,81,91,105,119,133,147,161,204) | *fla* |
| unk_1486 | (105,77) | *reg* |
| unk_1493 | (41,55,79,91,105,119,133,147,161,204) | *fla* |
| unk_1513 | (43,61,69,81,99,127,159) | *cal* |
| unk_1516 | (41,55,79,81,91,105,119,133,147,161,189,204) | *fla* |
| unk_1522 | (40,41,55,69,79,81,91,105,120,133,161,189,204) | *fla* |
| unk_1533 | (41,55,79,81,91,105,119,133,148,161,204) | *fla, bar* |
| unk_1535 | (43,55,69,81,93,105,119,133,149,161,189,207,222) | *fla* |
| unk_1539a | (41,55,77,81,91,105,119,134,161,189,204) | *fla* |
| unk_1539b | (41,44,69,91,115,142,157) | *fas, cal* |
| unk_1568 | (40,55,69,88,96,123,138,163,180,226) | *mic* |
| unk_1590 | (170,198,155) | *cal* |
| unk_1598 | (41,43,55,79,93,107,135,204) | *fas* |
| unk_1610 | (43,71,123,138) | *cal, mic* |
| unk_1624 | (41,43,55,69,79,93,107,135,204) | *fas* |
| unk_1638 | (40,5569,81,95,105,119,133,147,161,179,204) | *bar* |
| unk_1649 | (69,105,123,138) | *cal* |
| unk_1671 | (41,55,67,81,96,110,124,138,152,236) | *cali* |
| unk_1738 | (43,55,79,91,107,121,136,145,163,178) | *fla* |
| unk_1770 | (41,55,69,91,105,119,133,147,175,190,218) | *bar* |
| unk_1781 | (181,212) | *col* |

**Table S5. Global summary of total significantly differentially expressed transcripts (padj<0.05, ∣log2FC∣>1) across all treatment comparisons in *C. henryi*.**

| **Treatment Comparison** *(vs Control)* | **Tissue** | **Total DETs** | **Upregulated  (n, %)** | **Downregulated (n, %)** |
| --- | --- | --- | --- | --- |
| Removal | Labellum | 202 | 55 (27%) | 147 (73%) |
| Deposition | Labellum | 119 | 72 (61%) | 47 (39%) |
| Deposition+Removal | Labellum | 160 | 94 (59%) | 66 (41%) |
| Removal | Staminode | 217 | 90 (41%) | 127 (59%) |
| Deposition | Staminode | 128 | 81 (63%) | 47 (37%) |
| Deposition+Removal | Staminode | 176 | 134 (76%) | 42 (24%) |
| Removal | Petals | 819 | 583 (71%) | 236 (29%) |
| Deposition | Petals | 1010 | 670 (66%) | 340 (34%) |
| Deposition+Removal | Petals | 989 | 676 (68%) | 313 (32%) |

**Table S6. Differentially expressed transcripts (DETs) of candidate genes involved in terpene and fatty acid biosynthetic pathways in *C. henryi.*** Expression levels under different pollination treatments are shown relative to the control. Tissue abbreviations: S, staminode; L, labellum; P, (lateral) petals.

| **DET** | **Gene** | **Treatment** | **Tissue** | **Log2FC** | **P-value** | **Padj-value** |
| --- | --- | --- | --- | --- | --- | --- |
| trinity_dn10003_c0_g2_i11 | ***HMGCR3*** | Removal | *S* | 16,9103683 | 2,77E-05 | 0,01118986 |
|  |  | Deposition | *S* | 18,5497128 | 7,09E-07 | 0,00073971 |
|  |  | Dep + Rem | *L* | 18,4141619 | 6,49E-07 | 0,00062746 |
|  |  | Removal | *P* | 20,4622329 | 1,80E-07 | 4,62E-05 |
|  |  | Deposition | *P* | 16,6470897 | 8,85E-06 | 0,00095761 |
|  |  | Dep + Rem | *P* | 18,3143676 | 6,56E-07 | 0,00011196 |
| trinity_dn15849_c0_g1_i9 | ***RABGAP22*** | Removal | *S* | 18,4808825 | 1,84E-08 | 2,10E-05 |
|  |  | Dep + Rem | *S* | 20,0166582 | 6,34E-11 | 1,72E-07 |
|  |  | Removal | *P* | 16,1845637 | 8,46E-07 | 0,00017199 |
|  |  | Deposition | *P* | 17,4248311 | 2,68E-08 | 6,29E-06 |
|  |  | Dep + Rem | *P* | 17,9936999 | 4,53E-09 | 1,36E-06 |
| trinity_dn10588_c0_g1_i2 | ***ICL*** | Dep + Rem | *S* | 11,9682305 | 8,64E-05 | 0,03252777 |
|  |  | Removal | *P* | 17,7732028 | 1,02E-08 | 3,68E-06 |
|  |  | Deposition | *P* | 15,8260911 | 1,10E-07 | 2,14E-05 |
|  |  | Dep + Rem | *P* | 18,430522 | 2,30E-10 | 9,67E-08 |
| trinity_dn2196_c0_g1_i11 | ***ACX1_1*** | Removal | *P* | 6,42942083 | 1,69E-07 | 4,40E-05 |
|  |  | Deposition | *P* | 5,75539619 | 1,68E-06 | 0,00022902 |
|  |  | Dep + Rem | *P* | 6,23926103 | 1,90E-07 | 3,80E-05 |
| trinity_dn2196_c0_g1_i7 | ***ACX1_2*** | Dep + Rem | *P* | 4,5845751 | 0,00041114 | 0,02575175 |
| trinity_dn1890_c0_g1_i13 | ***ACX3*** | Removal | *P* | 19,3058915 | 2,39E-06 | 0,00041878 |
|  |  | Deposition | *P* | 18,082552 | 3,48E-06 | 0,00043446 |
|  |  | Dep + Rem | *P* | 17,7246165 | 3,32E-06 | 0,00047285 |
| trinity_dn3963_c0_g1_i6 | ***KCS2*** | Deposition | *P* | -4,43099713 | 0,00032896 | 0,02068241 |
| trinity_dn3963_c0_g1_i1 | ***KCS20*** | Deposition | *S* | -15,236395 | 1,95E-06 | 0,00167223 |
|  |  | Deposition | *L* | -14,7321524 | 4,27E-06 | 0,00405204 |
| trinity_dn81_c0_g1_i12 | ***ENR*** | Dep + Rem | *S* | -6,57751542 | 5,73E-05 | 0,0223338 |
|  |  | Removal | *P* | 7,52993705 | 1,46E-05 | 0,0019726 |
|  |  | Deposition | *P* | 6,50561494 | 0,00012595 | 0,00929916 |
|  |  | Dep + Rem | *P* | 5,73087045 | 0,00097587 | 0,04957011 |
| trinity_dn1882_c0_g1_i3 | ***GlyMFP-α*** | Deposition | *P* | 6,09254458 | 0,00023658 | 0,01572459 |
| trinity_dn4014_c0_g2_i1 | ***PDH-E1-α3*** | Deposition | *P* | 2,4068271 | 0,00078459 | 0,04078701 |
|  |  | Dep + Rem | *P* | 2,55199464 | 0,00046498 | 0,02830842 |
| trinity_dn546_c1_g1_i29 | ***BCCP1_1*** | Removal | *P* | -22,6517197 | 7,07E-10 | 3,81E-07 |
|  |  | Dep + Rem | *P* | -23,0193734 | 4,01E-12 | 2,47E-09 |
| trinity_dn546_c1_g1_i36 | ***BCCP1_2*** | Removal | *P* | 3,09483117 | 2,47E-08 | 8,10E-06 |
|  |  | Deposition | *P* | 2,82656103 | 1,17E-08 | 3,05E-06 |
|  |  | Dep + Rem | *P* | 2,46466216 | 9,51E-06 | 0,00116734 |
| trinity_dn401_c0_g1_i10 | ***PMPCB*** | Dep + Rem | *L* | 7,87657535 | 4,91E-05 | 0,02623691 |
| trinity_dn12045_c0_g1_i66 | ***ZC3H55*** | Removal | *S* | 16,0899652 | 1,92E-06 | 0,00117017 |
|  |  | Deposition | *S* | 15,6988061 | 1,12E-06 | 0,00105325 |
|  |  | Dep + Rem | *S* | 20,6537403 | 4,26E-11 | 1,27E-07 |
| trinity_dn19872_c0_g1_i15 | ***TNT1-94 Pol*** | Deposition | *L* | 26,0231823 | 3,49E-06 | 0,00336166 |
| trinity_dn813_c3_g1_i10 | ***BADC*** | Removal | *L* | -21,5947959 | 3,92E-06 | 0,00221434 |
|  |  | Removal | *P* | -24,3923668 | 1,79E-07 | 4,62E-05 |
|  |  | Dep + Rem | *P* | -24,6600845 | 6,47E-09 | 1,87E-06 |
| trinity_dn5020_c0_g1_i2 | ***FAD4*** | Deposition | *P* | -28,0249982 | 3,42E-08 | 7,71E-06 |
| trinity_dn5966_c0_g1_i1 | ***PKA*** | Removal | *P* | -24,9306557 | 1,57E-13 | 2,64E-10 |
|  |  | Deposition | *P* | -26,1586866 | 3,57E-17 | 1,27E-13 |
| trinity_dn10597_c0_g1_i44 | ***LACS6*** | Removal | *P* | -23,1895574 | 7,17E-17 | 3,62E-13 |
|  |  | Deposition | *P* | -24,6308369 | 1,72E-22 | 4,31E-18 |
|  |  | Dep + Rem | *P* | -24,3613039 | 6,36E-23 | 6,36E-19 |
